# Microglia support regeneration of rod photoreceptors in larval zebrafish retina through Nrf2-independent mechanisms

**DOI:** 10.64898/2026.10.22.761077

**Authors:** Michael Morales, Diana M Mitchell

## Abstract

The molecular and cellular components of successful regeneration of complex tissues in zebrafish are a major area of investigation. Previous work by our lab indicates a redox function for retinal microglia in the first 48 hours following acute injury to rod photoreceptors in the larval zebrafish retina. In addition, recent literature suggests that microglia provide signals that impact the regenerative response in the damaged retina. Here, we investigate the requirement of microglia and the redox-responsive transcription factor Nrf2 in the spontaneous regeneration of rod photoreceptors in the larval zebrafish retina. We used *irf8* mutants which are deficient in microglia, and mutant zebrafish lacking a functional copy of the gene *nfe2l2a*, which encodes a zebrafish ortholog of Nfe2l2/Nrf2. Our results indicate that microglia support the production of regenerated rods following their ablation. However, although detection of reactive oxygen species was increased in *nfe2l2a* mutant microglia following rod ablation, the regeneration of rods was not impacted in *nfe2l2a* mutants. Collectively, our results indicate a role for microglia in supporting the production of regenerated retinal neurons while also indicating that Nrf2-independent mechanisms can likely provide redox functions during rod regeneration.

## Introduction

The regeneration of retinal neurons in zebrafish has been studied with the goal of translating such knowledge to rescue vision loss in humans. This is because the zebrafish retina can spontaneously regenerate neurons following damage, a capability not shared by humans. The Müller glia have been identified as a source of neuronal progenitors in the zebrafish retina following acute retinal damage (1–5). In addition to continued work aimed to understand how Müller glia transition from homeostasis to a regenerative response, the function of microglial cells (the resident macrophages of the central nervous system) in facilitating or supporting the process of regeneration has recently gained strong interest. The quest to understand microglial functions in retinal regeneration has produced a body of work aimed to understand the requirement and pertinent mechanisms for microglia in supporting regeneration of the tissue (6–15). For example, inflammatory signals from microglia may be a factor that induces re-entry of Müller glia into the cell cycle to generate neuronal progenitors (12, 15–17). Work from our lab and others strongly suggests that an inflammatory microglial phenotype impacts the survival of regenerated retinal neurons (7, 17), though a specific requirement for microglia presence to support survival of regenerated neurons remains unclear (15). Overall, microglial functions towards differentiation, survival, and re-establishment of synaptic networks of regenerated neurons currently remain poorly understood.

Redox balance may be an important component of repair and regeneration following retinal damage (6, 18, 19). A group of molecules collectively called reactive oxygen species (ROS) are key players in redox biology. These molecules exist both as radicals, such as superoxide (O2^-^) and hydroxyl (OH•), or as non-radicals such as hydrogen peroxide (H2O2), and possess varying propensities to react with other biomolecules (20). Often, ROS are considered damaging molecules because they can cause oxidative damage, but it is also known that they have important roles in modulating the activity of signaling pathways involved with cell proliferation and differentiation (20, 21), the hypoxia response, autophagy (22, 23), and inflammation (24–27). In order to mitigate the risk of oxidative damage, cells possess a robust antioxidant system consisting of enzymes and other molecules. We previously showed that microglia express several enzymes important for redox biology (6, 19) and as scavengers and phagocytes, microglia are well-suited for both the generation and detoxification of oxidants (28). Indeed, we identified microglia as an important factor towards restoration of redox balance after retinal photoreceptor damage (18).

The transcription factor nuclear factor erythroid 2-related factor 2 (Nfe2l2, also known as Nrf2) is a master transcriptional regulator of the antioxidant response. (29)Nrf2 is normally bound in the cytosol to Kelch-like ECH-associated Protein 1 (KEAP1) and is released for nuclear translocation upon oxidation of cysteine residues on KEAP1 (29), thus rendering it redox sensitive. When active and localized to the nucleus, Nrf2 binds to a regulatory sequence known as the antioxidant response element (ARE) in its target genes (29). We previously profiled transcripts in zebrafish microglia at 7 days following ouabain-induced damage, a timepoint when the Müller glia have generated actively dividing neuronal progenitor cells (6). Included in this “regeneration-associated transcriptome” of zebrafish microglia was expression of the Nrf2 ortholog *nfe2l2a* and several other antioxidant genes (6), suggesting that antioxidant function of microglia is involved in retinal regeneration. In addition, we detected increased expression of *nfe2l2a* in microglia-deficient (*irf8-/-*) larval zebrafish eyes (18), suggesting that microglial antioxidant functions are also an important element of retinal homeostasis. In the same study, which used a drug-inducible system for targeted rod photoreceptor ablation (30), we found that an oxidative load is generated in the dying rods (18). This load was undertaken and mitigated by microglia during the early damage response, and microglia were required to restore the redox homeostasis of the retina (18). Other studies indicate a connection between redox balance and retinal regeneration. A study by *Boyd and Hyde* found that iron, a highly redox-active transition metal, and glutathione peroxidase 4b are important in promoting Müller glia proliferation in zebrafish (31). Additionally, a study by *Emmerich et al.*, found that the CRISPR-mediated knockout of thioredoxin was found to enhance the regeneration of rod photoreceptors in larval zebrafish (32). Collectively, these papers suggest a link between redox balance, microglia, and retinal regeneration in zebrafish. In other tissues, a similar link is also suggested. In the brain, it was shown that stabilizing the activity of Nrf2 within microglia facilitated tissue recovery from white matter ischemia and remyelination of neurons (33).

In this study, we investigate the requirements for microglia and Nrf2 function in the regenerative response following rod photoreceptor damage in larval zebrafish retina. We hypothesized that Nrf2 is important for the viability and activation of microglia, and that microglia and Nrf2 are important factors for the transition from damage to regeneration response. As tools to test this hypothesis, we used *rho:nfsB-eGFP^nt19* (30) transgenic zebrafish to ablate rods upon exposure to the drug Metronidazole (MTZ) and trigger their regeneration, and the *irf8^st95* (34) mutation for microglial deficiency (18). Additionally, we incorporated a loss-of-function mutant of *nfe2l2a* (*nfe2l2a^w211*) (35) to determine if Nrf2 function is required for the regeneration of rod photoreceptors. We found that the production of regenerated rods was compromised in the absence of microglia, which was likely mitigated by additional progenitor proliferation and/or rod production in *irf8* mutants. While *nfe2l2a* mutants had increased ROS loads in microglia responding to rod cell death, surprisingly, retinal microglia in *nfe2l2a* mutants did not show significantly altered morphology or viability. Further, *nfe2l2a* mutants did not show significant impacts to rod regeneration. Our results indicate that microglia support the production of regenerated rod photoreceptors, either for their proper differentiation and/or their survival, but that redox functions of microglia in the regenerating retina can be executed in an Nrf2-independent manner.

## Results

### Rod ablation in microglia-sufficient and microglia-deficient larval retinas primarily results in proliferation in the outer nuclear layer

We aimed to determine if microglia, possibly through their function to restore redox balance, were required for the regeneration of rod photoreceptors. As in our previous paper (18), we used *rho:nfsB-eGFP* (*^nt19*) transgenic larval zebrafish, which allow rod ablation upon exposure to MTZ (30). This is because nfsB encodes a nitroreductase enzyme (NfsB) that converts MTZ to a toxic product, leading to death of nfsB-expressing cells, which in this system are rod photoreceptors. In addition, our previous paper showed that coupled to their death, the nfsB-expressing rods exposed to MTZ accumulated ROS (18). We followed the same 48-hour treatment procedure that we used previously to induce rod cell death (18): larval *rho:nfsB-eGFP* were exposed to DMSO (vehicle control) or MTZ for 48 hours, then then washed and transferred to fresh water for recovery. In these experiments, we also included the *irf8*^st95 mutation (34) and collected samples from *irf8*+/-(microglia sufficient) and *irf8*-/-(microglia deficient) siblings. Samples were collected at 24-hour intervals beginning at 48 hours post-treatment (hpt, immediately following the end of DMSO/MTZ treatment) and through 120 hpt (three days of recovery after drug washout). For clarity, the term “hours post-treatment” is relative to the initiation of treatment. We previously demonstrated ablation of rods in larval *rho:nfsB-eGFP* retinas by 48 hours following MTZ treatment (18). Consistent with rod ablation by MTZ exposure, we again found a drastic loss of GFP+ rod photoreceptors by 48 hpt and this was evident in both *irf8* genotypes (Figure 1A-A’’’ and Supplemental Figure 1A-A’’’).

**Figure 1:**
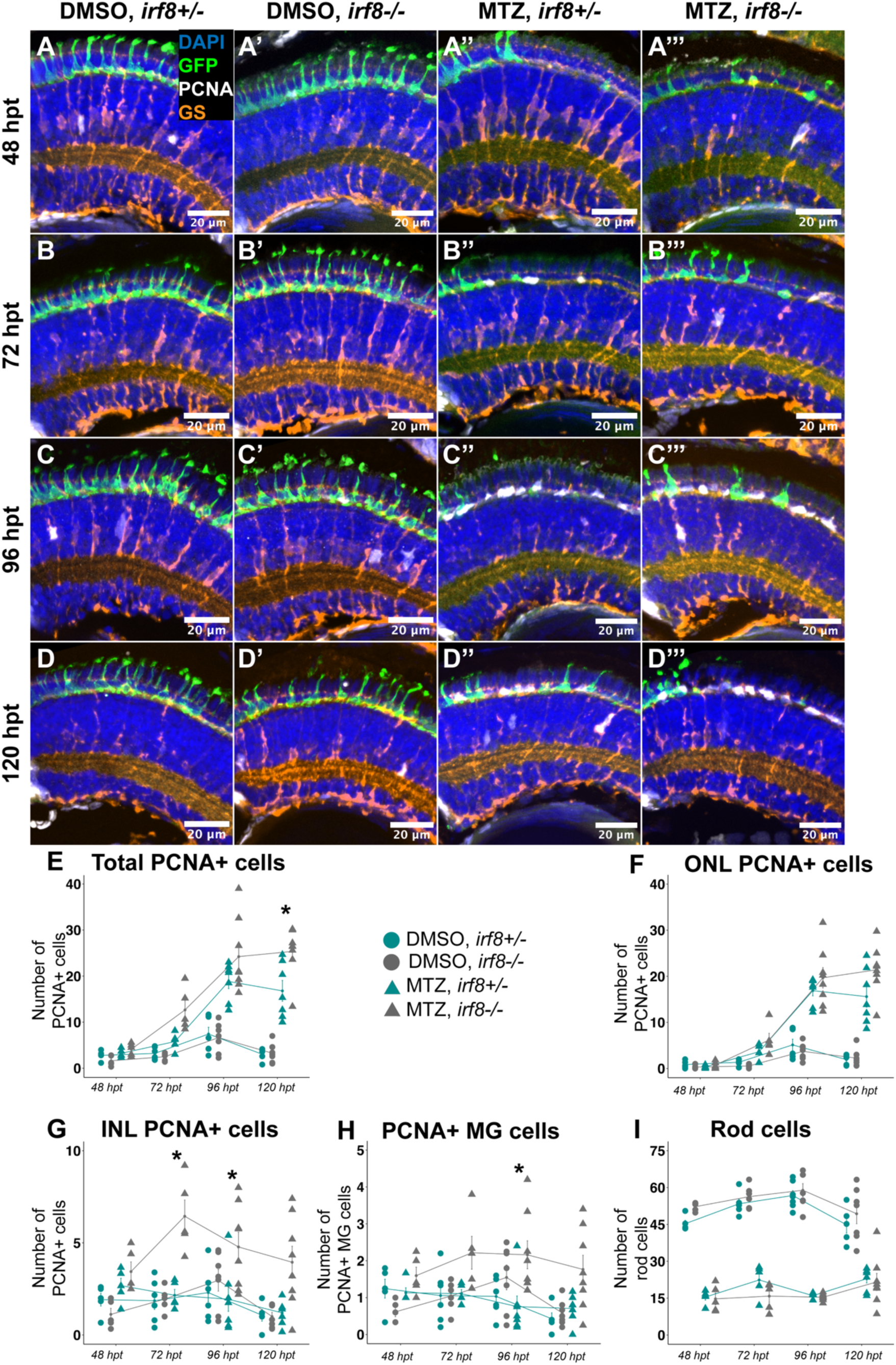
Cell proliferation following rod ablation in *irf8*+/- and *irf8-/-*retinas. A-D’’’: Representative images of dorsal region of retinas from retinal cryosections showing DAPI (blue), GFP+ rods (green), PCNA (white), and GS (orange) in DMSO (A-D, A’-D’) and MTZ-treated (A’’-D’’, A’’’-D’’’) and either *irf8+/-*(A-D, A’’-D’’) or *irf8-/-*(A’-D’, A’’’-D’’’). Top row shows samples at 48 hpt (A-A’’’); second row shows samples at 72 hpt (B-B’’’); third row shows samples at 96 hpt (C-C’’’), bottom row shows samples at 120 hpt (D-D’’’). E-I: Average number of total PCNA+ cells (E), ONL PCNA+ cells (F), and INL PCNA+ cells (G), PCNA+ Müller glia cells (H), and Rod cells per retinal cryosection over time from 48-120 hpt. Each point represents the average value of cells from retinal cryosections; 5-8 eyes; 3-5 larvae. Cyan circles denote DMSO *irf8+/-*; gray circles denote DMSO *irf8-/-*; cyan triangles denote MTZ *irf8+/-*; gray triangles denote MTZ *irf8-/-*. *Indicates significant difference between MTZ *irf8+/-* and MTZ *irf8-/-*where p<0.05, two-way ANOVA with Tukey’s HSD post-hoc test for multiple comparisons.

Following neuronal damage in the teleost retina, a proliferating population of multipotent progenitor cells originates from the Müller glia, which are called Müller glia-derived progenitor cells (MGPC) (2–5, 36–38). In addition, a population of unipotent rod progenitors known as rod precursor cells exist in the zebrafish retina that amplify proliferation for the specific replacement of rods (39, 40). In adult *rho:nfsB-eGFP* (*^nt19*) retinas, the ablation of rods is sufficient to induce the production of MGPC (30); however, the proliferative profile in larval retinas of this transgenic line has not been shown. We therefore used PCNA staining to visualize proliferating progenitors following MTZ exposure and washout. Retinal layers were identified by DAPI staining, and we also included Glutamine Synthetase (GS) staining to identify the Müller glia (MG) including those that had entered asymmetric cell division (5, 41). We performed immunofluorescent staining on retinal cryosections of *rho:nfs*B-eGFP larvae of *irf8+/-*(microglia-sufficient) and *irf8-/-*(microglia-deficient) genotypes collected at successive 24-hour timepoints beginning at the cessation of DMSO/MTZ exposure (48 hpt). We assessed PCNA staining in the INL and ONL of retinal cryosections, with INL PCNA serving as a proxy of MGPCs, and PCNA staining in the ONL serving as a proxy measure of the amplification of MGPC, which migrate apically (2, 42), and rod precursor cells, which also localize to the ONL (2, 39). DMSO-treated (undamaged) retinas had very few PCNA+ cells in all layers throughout the time course, while proliferation in MTZ-treated groups increased at 96 hpt (Figure 1 A-E). (30). We also noted that PCNA+ cell abundance following MTZ treatment was visibly more noticeable in the dorsal region of the retina (Supplemental Figure 1).

In microglia sufficient *irf8*+/-samples, PCNA+ cells were primarily detected in the ONL (Figure 1 F) following MTZ treatment, while PCNA+ cell numbers in the INL remained similar to DMSO controls (Figure 1 G). In *irf8*-/-retinas compared to *irf8*+/-following rod ablation, there were slightly increased numbers of PCNA+ cells in the ONL at 72 and 96 hpt, and the numbers of PCNA+GS+ Müller glia were slightly elevated at 96 hpt. However, it is worth noting that, overall, the numbers of PCNA+GS+ cells (considered to be dividing Müller glia) were very small, with a group average of less than 3 PCNA+ GS+ cells per section (Figure 1 H). Total PCNA+ numbers were elevated in *irf8-/-*retinas compared to *irf8+/-*retinas at 120 hpt following MTZ treatment (Figure 1 E). Quantification of GFP+ rod numbers in MTZ treated samples compared to DMSO controls showed that rod numbers remained reduced and were not yet restored to levels of undamaged conditions by 120 hpt (Fig 1 I). The number of GFP+ rods was similar in both *irf8* genotypes of both DMSO and MTZ treated samples across the timepoints examined (Fig 1 I). These results indicate that the ONL is the primary site of proliferation in response to rod ablation, and Müller glia cell proliferation was not strongly induced in *rho:*nfsB-eGFP larval retina following MTZ-mediated rod ablation. Further, modest differences in PCNA abundance in MTZ-treated *irf8+/-* and *irf8-/-*retinas, suggested slightly elevated levels of cell proliferation in microglia-deficient samples, though this may be transitory and unsynchronized.

### Microglia-deficiency impacts the outcome of rod photoreceptor regeneration

To more directly analyze regeneration of rods in larval retinas in both microglia-sufficient and microglia-deficient conditions, we used 5-ethanyl 2’-deoxyuridine (EdU) incorporation to label rods produced from proliferating progenitors following rod ablation. We exposed *rho:nfsB-eGFP* larvae of both *irf8* genotypes to EdU via immersion in fish water from 48-72 hpt. We selected this timing due to the presence of PCNA+ cells in the INL temporally preceding the more pronounced proliferation in the ONL at 96 hpt, suggesting initial stages of the regenerative response (Figure 1). Further, this EdU exposure timing would allow us to track rods produced from both MG and rod precursor niches. We collected samples at 216 hpt (9 days after the onset of MTZ treatment and 7 days after MTZ washout), which allowed larvae 7 days of recovery and regeneration. To quantify regenerated EdU+ rods, we counted EdU+ GFP+ rods that were located in between the EdU+ ciliary marginal zone (CMZ), visible as a band of strongly labeled EdU+ cells near the periphery of the retinal sections (Figure 2). This allowed us to identify EdU+ rods in the area of the retina that experienced rod cell death during treatment as opposed to regions that developed afterwards from the CMZ. In DMSO-treated retinas, we did not detect significant numbers of EdU+ rods, and overall rod cell numbers were similar in both *irf8* genotypes, suggesting that differentiation of rods during retinal development is not affected by microglia deficiency at this age (Figure 2 A, A’, F-K). MTZ-treated retinas had significantly increased numbers of EdU+ rods (Figure 2 A’’,A’’’, F-K), consistent with, and as expected for, rod regeneration following their ablation and MTZ washout. The EdU+ rods were localized predominantly on the dorsal side of the retina (Fig 2 G-H), consistent with localization of PCNA+ progenitors described above (Supplemental Figure 1). Interestingly, we found that the number of EdU+ rods were reduced in MTZ-treated *irf8-/-*retinas compared to *irf8+/-*(Figure 2 D-E’, F-H). However, despite the fewer numbers of EdU+ rods in *irf8-/-*retinas, the overall GFP+ rod cell numbers were not different between *irf8+/-* and *irf8*-/-retinas (Figure 2 I-K). This discrepancy between EdU+ rod and total rod numbers may indicate an impact to survival of regenerated rods in *irf8* mutants or alternatively, altered timing of rod differentiation relative to EdU exposure.

**Figure 2:**
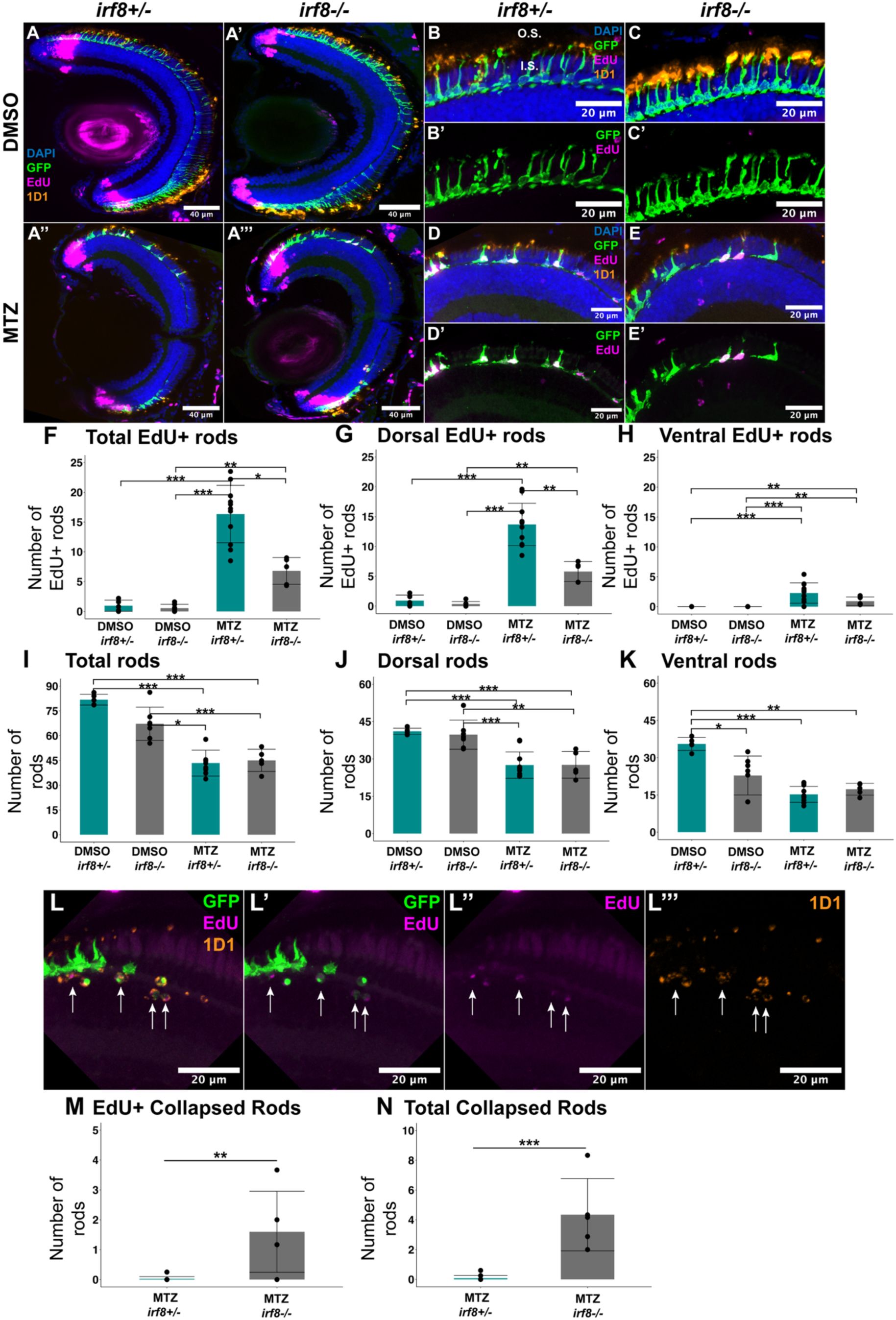
Analysis of regenerated rods using EdU at 216 hpt. **A-A’’’:** Representative images of retinal cryosections at 216 hpt (9 days post-injury) from DMSO-(top row) and MTZ-treated retinas (bottom row) with either *irf8+/-*(left-hand column) or *irf8-/-*(right-hand column) genotype. Images show DAPI (blue), GFP rods (green), EdU (magenta), and 1D1 (rhodopsin, orange). **B-E’:** Dorsal regions of retinal cryosections at 216 hpt showing DAPI (blue), GFP rods (green), EdU (magenta), and 1D1 (rhodopsin, orange) (four-channel images B, C, D, E) or dual-channel images showing just GFP rods and EdU (B’, C’, D’, E’). Upper row shows DMSO-treated retinas with either *irf8+/-*(B, B’) or *irf8-/-*(C, C’) genotype. Lower row shows MTZ-treated (regenerating) retinas with either irf8+/-(D, D’) or irf8-/-(E, E’) genotype. O.S. and I.S. in B indicate outer- and inner segment regions, respectively. **F-H:** Counts of EdU+ rods per retinal cryosection in the total retinal section (F), dorsal (G), and ventral (H) retina. Bars represent group means +/- SD, and each dot represents the average count per eye. **I-K:** Counts of total rods per retinal cryosection in the total retinal section (I), dorsal (J), and ventral (K) retina. Bars represent group means +/-SD, and each dot represents the average count per eye. L-L’’’: Representative sample of a region of collapsed rods in MTZ-treated *irf8-/-*retinas at 216 hpt. Images show either GFP (green), EdU (magenta), and 1D1 (orange) channels (L), or just GFP and EdU (L’), EdU alone (L’’), or 1D1 alone (L’’’). Arrows indicate collapsed rods. M-N: Numbers of collapsed EdU+ rods (M) and total collapsed rods (N) in regenerating retinas. Bars represent mean +/-SD and dots represent average number of collapsed rods per eye. n=5-10 eyes (3-6 larvae). *p<0.05, ** p<0.01, ***p<0.001, two-way ANOVA with Tukey’s HSD except Conover test with Bonferroni correction (H) and Mann-Whitney U-test (M, N).

We further analyzed the features and morphology of the EdU+ GFP+ rods in the two *irf8* genotypes. In addition to the *rho*-driven GFP reporter to visualize rod cell bodies, we used the antibody 1D1 to label the rod photopigment rhodopsin (43), which is normally localized to the outer segments of rods (44), as shown readily in our control groups (Figure 2 B,C, denoted “O.S.”). However, in *irf8*-/-regenerating retinal tissue, we observed the presence of abnormally shaped rod cells: these rods had GFP+ cell bodies with a circular, collapsed shape and lacked visible inner/outer segment extension with rhodopsin staining around the periphery of the circular GFP+ center (Figure 2L-L’’, M-N). We defined these as “collapsed” GFP+ rods and noted that they sometimes appeared in clusters in the ventral region of *irf8*-/-retinal sections (Figure 2 L-L’’’, M-N). GFP+ collapsed rods were essentially absent from *irf8+/-*retinal samples where rods instead showed overall typical morphology and rhodopsin localization to the O.S. (Figure 2 M-N, Supplemental Figure 2 A). Several of the collapsed GFP+ rods in *irf8* mutants also stained positive for EdU (Figure 2 L-L’’, arrows, M-N) indicating that they were derived from proliferating progenitors between 48 and 72 hpt. Rhodopsin staining was detected within the collapsed GFP+ cell bodies (Figure 2 L, L’’’, arrows), which may indicate that these GFP+ rods had either formed outer segments then collapsed, experienced rhodopsin mis-localization, and/or experienced slow or abnormal development. Such morphological abnormalities suggest that rather than merely altered timing of rod differentiation, the proper differentiation or survival of regenerated rods is likely compromised in *irf8-/-*retinas.

To further address these findings with additional temporal resolution, we examined samples from each group that had undergone the same EdU exposure window following rod ablation but were collected 3 days earlier at 144 hpt (6 days post-injury, 4 days recovery following MTZ-washout, Figure 3). Once again, we did not find significant numbers of EdU+ rod cells in DMSO-treated retinas (Figure 3 A-A’, B-C’, F-H), but we detected increased numbers of EdU+ rods in MTZ-treated retinas (Figure 3 A’’-A’’’, D-E’, F-H), and these were mostly dorsally localized (Figure 3 G). At this timepoint (144 hpt), there were no differences in the numbers of either EdU+ rods or in total rods between the two *irf8* genotypes (Figure 3 F-K), indicating that initial regeneration of rods (and EdU incorporation) proceeded similarly in both microglia-sufficient and -deficient retinas. However, when we analyzed the tissue for the presence of collapsed rods at 144 hpt, we again found such collapsed GFP+ rods in *irf8-/-*but not *irf8*+/-retinas (Figure 3 A’’’, arrows, L-L’’’). Additionally, many of these collapsed rods were also EdU+ (Figure 3 M-N). Again, collapsed rods were virtually absent from *irf8+/-*retinas (Figure 3 M-N, Supplemental Figure 2 C). Though MTZ may have some toxicity in microglia-deficient tissue, the collapsed rods in *irf8-/-*were detected at 144 and 216 hpt, 4 and 7 days after the removal of MTZ. Further, our previous findings using *rho*:nfsb-eGFP; *irf8-/-*mutants suggested that compensatory clearance activity in *irf8-/-*retinas is effective at clearing rods and other dying cells from the tissue by 72 hpt (18), so it is unlikely that even EdU-collapsed rods are remnants of the initial MTZ treatment. We also did not detect collapsed rods in the CMZ-adjacent regions of the retina which would have developed from the CMZ after MTZ-treatment, so impacts to rods appear to be confined to regions that previously experienced cell death rather than that which is newly developed from growth ant the peripheral margins.

**Figure 3:**
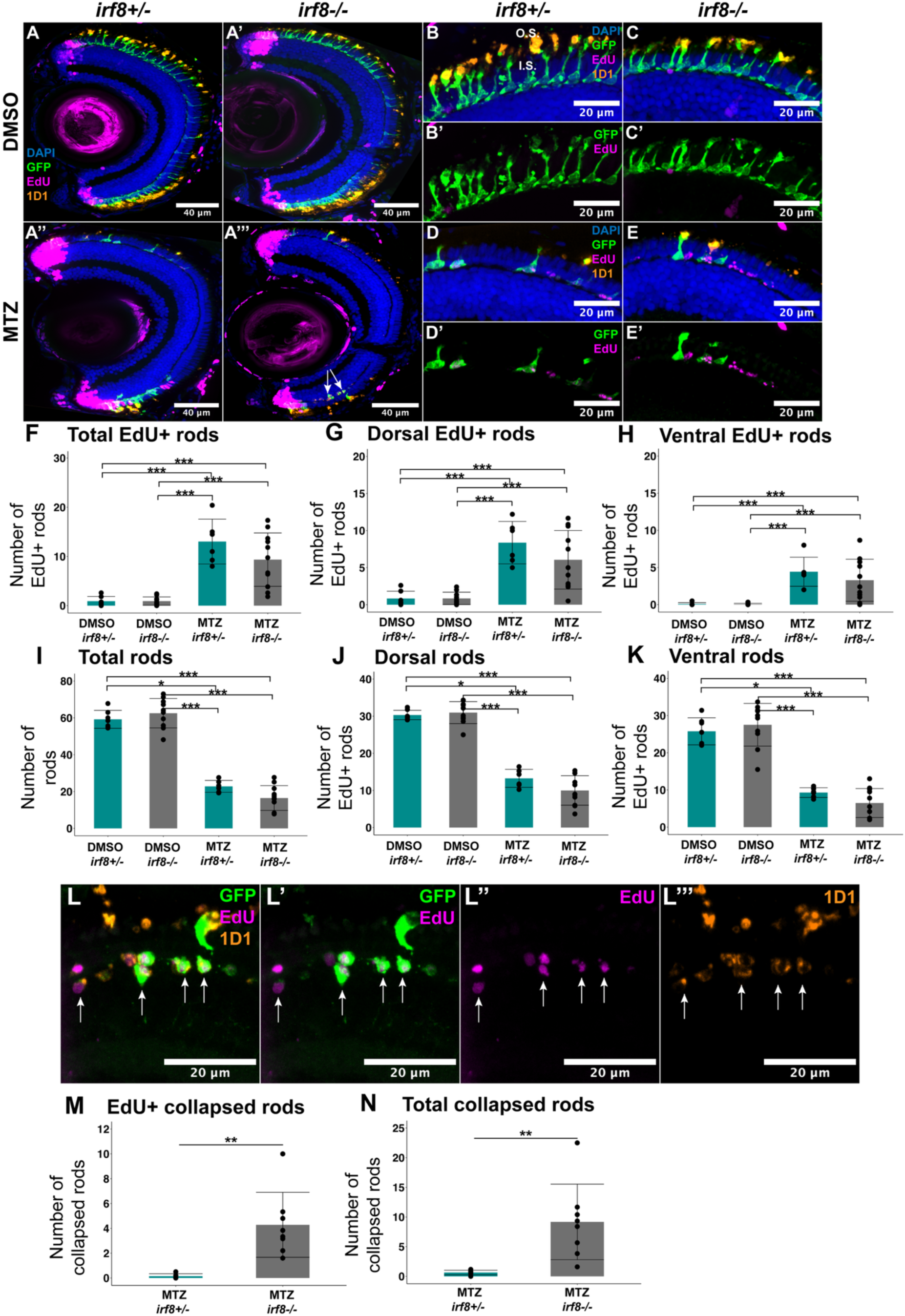
Analysis of regenerated rods using EdU at 144 hpt. **A-A’’’:** Representative images of retinal cryosections at 144 hpt (6 days post-injury) from DMSO-(top row) and MTZ-treated retinas (bottom row) with either *irf8+/-*(left-hand column) or *irf8-/-*(right-hand column) genotype. Images show DAPI (blue), GFP rods (green), EdU (magenta), and 1D1 (rhodopsin, orange). **B-E’:** Dorsal regions of retinal cryosections at 144 hpt showing DAPI (blue), GFP rods (green), EdU (magenta), and 1D1 (rhodopsin, orange) (four-channel images B, C, D, E) or dual-channel images showing just GFP rods and EdU (B’, C’, D’, E’). Upper row shows DMSO-treated retinas with either *irf8+/-*(B, B’) or *irf8-/-*(C, C’) genotype. Lower row shows MTZ-treated (regenerating) retinas with either irf8+/-(D, D’) or irf8-/-(E, E’) genotype. O.S. and I.S. in B indicate outer- and inner segment regions, respectively. **F-H:** Counts of EdU+ rods per retinal cryosection in the total retinal section (F), dorsal (G), and ventral (H) retina. Bars represent group means +/- SD, and each dot represents the average count per eye. **I-K:** Counts of total rods per retinal cryosection in the total retinal section (I), dorsal (J), and ventral (K) retina. **L-L’’’:** Representative sample of a region of collapsed rods in MTZ-treated *irf8-/-*retinas at 216 hpt. Images show either GFP (green), EdU (magenta), and 1D1 (orange) channels (L), or just GFP and EdU (L’), EdU alone (L’’), or 1D1 alone (L’’’). Arrows indicate collapsed rods. **M-N:** Numbers of collapsed EdU+ rods (M) and total collapsed rods (N) in regenerating retinas. Bars represent mean +/-SD and dots represent average number of collapsed rods per eye. n=5-10 eyes (3-5 larvae). *p<0.05, ** p<0.01, ***p<0.001, two-way ANOVA with Tukey’s HSD except Conover test with Bonferroni correction (H) and Mann-Whitney U-test (M, N).

We compared the numbers of EdU+ rods in MTZ-treated retinal sections between 144 and 216 hpt (Figure 4 A-C) and found that the number of EdU+ rods increased modestly from 144 hpt to 216 hpt for *irf8+/-*retinas, with a statistically significant increase occurring in the dorsal retina (Figure 4 B). This was not the case for *irf8-/-*retinas where such numbers did not increase over time (Figure 4 A-C). The total number of GFP+ rods remained similar in DMSO groups at both timepoints and increased significantly from 144 hpt to 216 hpt in MTZ-treated groups (regenerating retinas) of both genotypes (Figure 4 D). Collectively, this analysis further supports the interpretation that EdU incorporation into regenerated rods is initially similar between *irf8* genotypes, but the EdU+ regenerated rods are lost from microglia-deficient retinas over time. Interestingly, the total number of rods (EdU+ and EdU-) is similar in both *irf8* genotypes at 216 hpt, suggesting that additional production of rods may occur in microglia-deficient retinas, possibly to compensate for their lack of successful development or survival. When comparing across timepoints, we found slightly higher numbers of collapsed EdU+ rods in *irf8*-/-retinas at 144 vs 216 hpt (Figure 4 G-H), suggesting that these abnormal rods are cleared and/or do not persist over time (rather than cessation of their production, in which numbers would be similar across timepoints). Consistent with this, there are also EdU-GFP+ collapsed rods in *irf8*-/-retinas, suggesting that abnormal rods are still produced after EdU washout (Figure 4 N).

**Figure 4:**
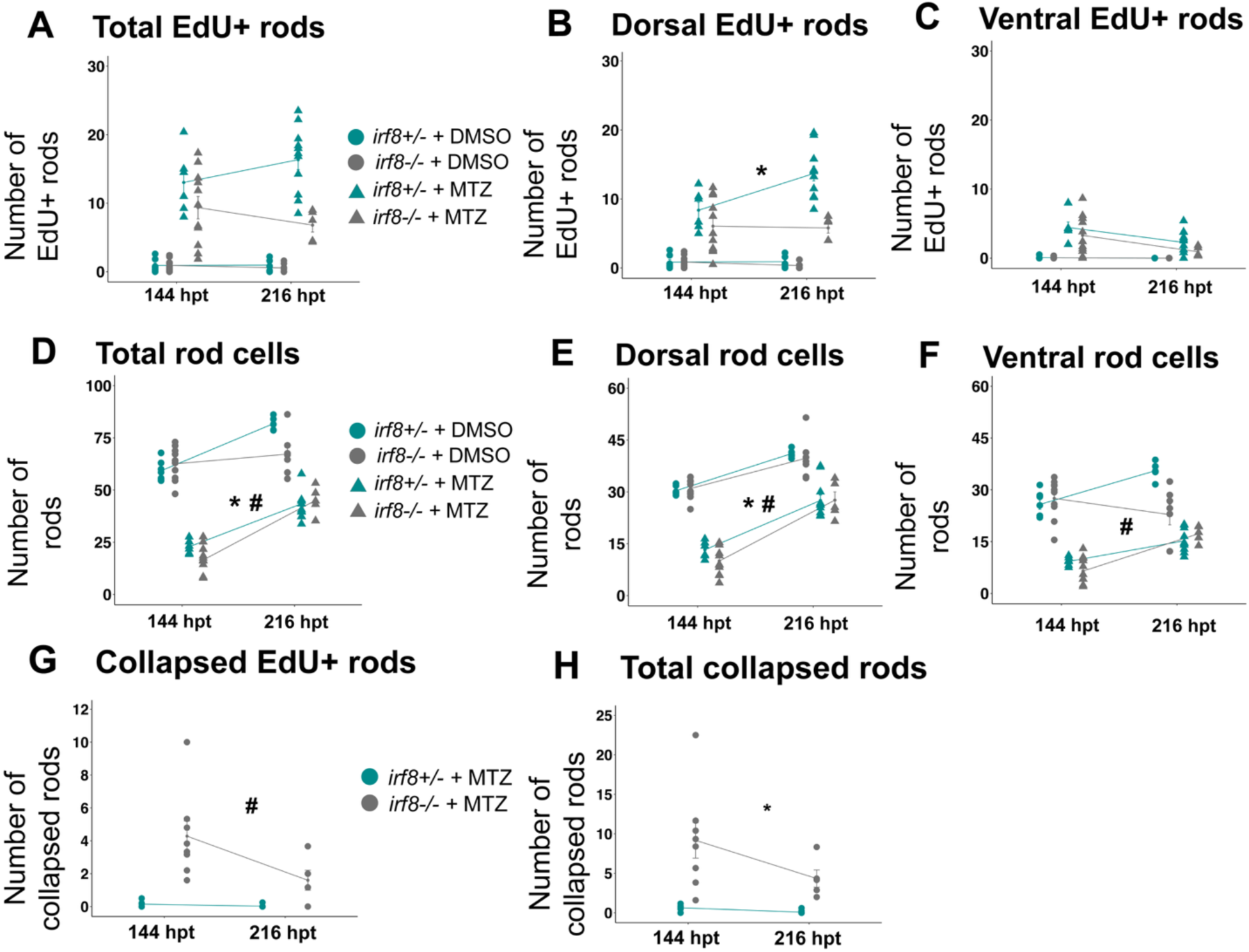
Comparison of EdU+ rods in regenerating retinas between 144- and 216 hpt. **A-C:** Counts of EdU+ rods per retinal cryosection in total retina (A), dorsal (B), and ventral (C) regions at both 144- and 216 hpt. **D-F:** Counts of total rods per retinal cryosection in total retina (D), dorsal (E), and ventral (F) regions at both 144- and 216 hpt. Each point represents the average number of rods (EdU or total, respectively) per cryosection. n=5-10 eyes, 3-6 larvae per group. Select comparisons annotated for statistical significance: * indicates statistical significance (p<0.05, estimated marginal means) for comparison between *irf8+/-*144 hpt and *irf8+/-*216 hpt. # indicates statistical significance (p<0.05, estimated marginal means) for comparison between *irf8-/-*144 hpt and *irf8 -/-*216 hpt. **G-H:** Collapsed EdU+ (G) and total (H) rods at 144- and 216 hpt. Dots represent average of technical replicates for an individual eye. *Indicates statistical significance (p<0.05, Conover test with Bonferroni correction) between *irf8+/-*regenerating retinas at 144 and 216 hpt. #indicates statistical significance (p<0.05, Conover test with Bonferroni correction) between *irf8-/-*regenerating retinas at 144- and 216 hpt.

### Loss of nrf2a function increases oxidative load on microglia but does not impact microglial response to rod ablation

Our previous work indicated that microglia express *nfe2l2a* (encoding Nrf2) and several of its target genes (6), and microglia support redox homeostasis of the retina following MTZ-mediated ablation of rods (18). In addition, our results above indicate that microglia support the production of regenerated rods. We therefore sought to determine if Nrf2-mediated mechanisms underlie the function of microglia following rod ablation. We crossed a loss-of-function mutation in the gene *nfe2l2a* (*nfe2l2a^w211*,(35)) with the *rho:nfsB-eGFP^nt19* transgenic line (18, 30) to generate *rho:nfsB-eGFP* Nrf2-sufficient and Nrf2-deficient larvae for MTZ-mediated rod ablation. Hereafter we refer to *nfe2l2a^w211 +/-*(heterozygotes, functional Nrf2) as *w211+/-*, and *nfe2l2a^w211 -/-*(homozygous mutants, Nrf2 loss-of-function) as *w211-/-*samples. We used the same DMSO/MTZ treatment schedules as previously (48 hours of drug exposure followed by washout).

We examined oxidative load in microglia responding to rod ablation (at 48 hpt) from both genotypes using CellRox™ to visualize ROS in larval zebrafish eyes/retinas (18) and quantifying (18)fluorescence intensity of CellRox™ signal (Figure 5 A-D’’). In both genotypes, rods were ablated upon MTZ treatment, evident from collapsed GFP+ rods visible in the ventral region of MTZ treated eyes (Figure 5 C’. D’, insets). Consistent with our previous findings (18), CellRox™ signal was minimal in DMSO treated samples (Figure 5 A’-B’’) but visible and co-localized to microglia in MTZ treated samples (Figure 5 C’-D’’). CellRox™ fluorescence intensity was increased in the microglia that were ventrally localized (rod-rich region) in response to MTZ-induced rod cell death compared to DSMO controls and this was the case for both *w211*+/- and *w211*-/-genotypes (Figure 5E). There was a slight decrease in CellRox™ fluorescence intensity in microglia from *w211-/-*DMSO-treated eyes compared to the *w211+/-*group (Figure 5 E). However, (18)CellRox™ fluorescence intensity was increased within microglia of *w211-/-*larval eyes compared to *w211*+/-eyes (Figure 5 E), indicating that loss of *nrf2a* increases ROS load on microglia upon rod damage.

**Figure 5:**
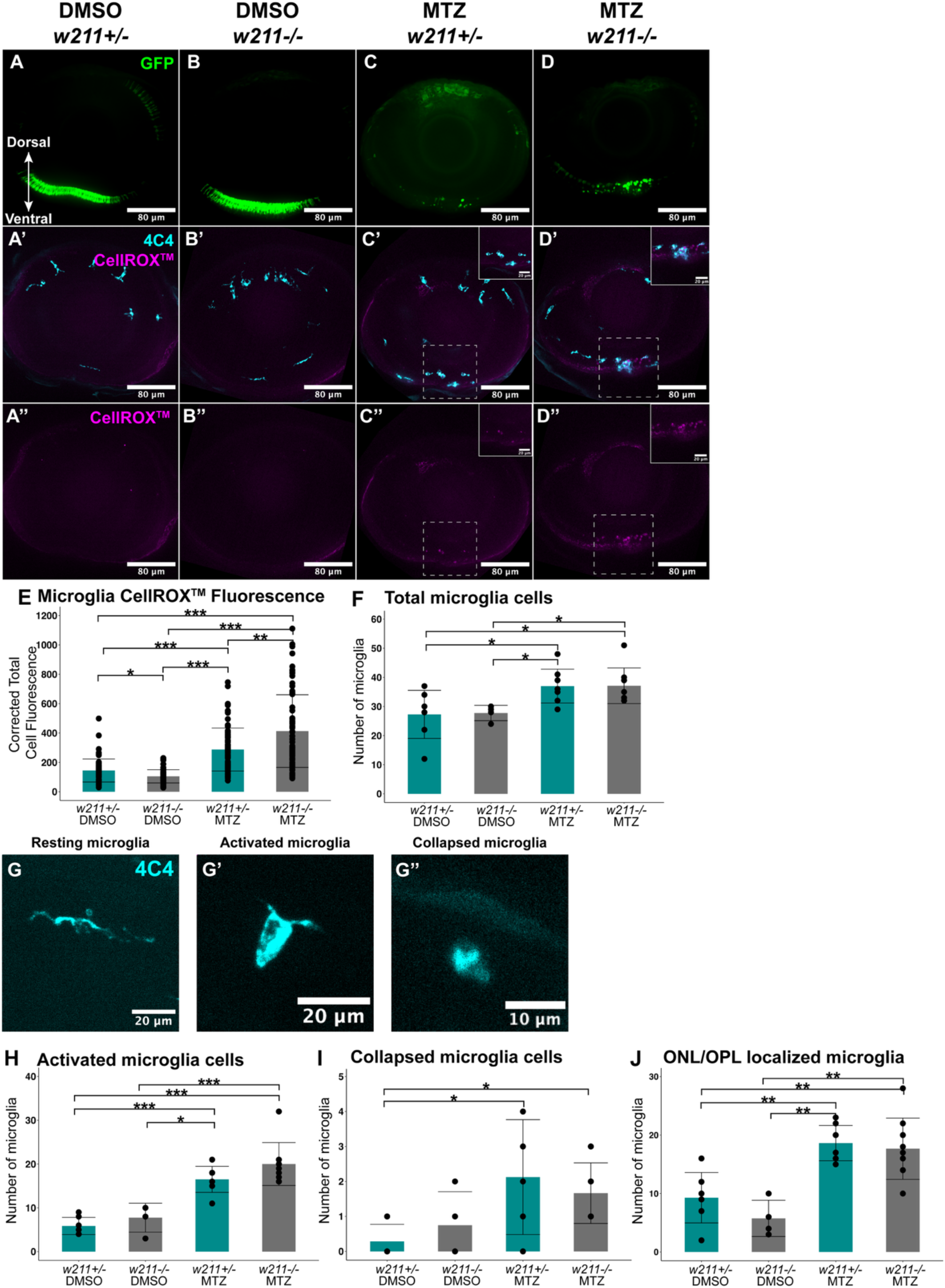
Analysis of microglial responses to rod death in the presence and absence of functional *nrf2*. **A-D’’:** Representative images of whole larval eyes treated with either DMSO (A-B’’) or MTZ (C-D’’) and either *w211+/-*(A-A’’, C-C’’) or *w211-/-*(B-B’’, D-D’’), scale bar 80 µm (insets 20 µm). Images in the left-hand column (A-D), show GFP+ rods in their respective groups. Signal is most intense in the ventral retina (arrows). Images in the middle column (A’-D’) show microglia (cyan) and CellRox™ signal in their respective groups. Microglia become more localized to the outer, ventral retina following rod ablation (C’, D’, insets). The right-hand column (A’’-D’’) shows single-channel images of CellRox™ signal. Signal becomes more intense and visible in the ventral retina following rod ablation (C’’, D’’, insets). **E:** Quantitative measurement of CellRox™ fluorescence intensity co-localized with microglia. Each datapoint represents fluorescence signal from single microglia; 4-9 eyes per group (3-5 larvae). **F:** Numbers of total microglia in whole larval eyes. Each datapoint represents the total number of microglia cells in a whole eye in the respective category, n=4-9 eyes, (3-5 larvae). **G-G’’:** Representative samples of defined morphological characteristics: resting, activated, and collapsed (G-G’’, respectively). **H-J:** Numbers of activated microglia (H), collapsed microglia (I), and ONL/OPL-localized microglia (J). Each datapoint represents the total number of microglia cells in a whole eye in the respective category, n=4-9 eyes, (3-5 larvae). *p<0.05, **p<0.01, ***p<0.001, two-way ANOVA with Tukey’s HSD post-hoc test for multiple comparisons. For clarity, only significant differences are shown in these graphs. Conover test (non-parametric) with Bonferroni correction for multiple comparisons in H.

To determine if the *nrf2* loss-of-function mutation had additional effects on microglia, we assessed numbers, morphological features, and localization of microglia in *w211+/-* and *w211-/-rho*:nfsB-eGFP larval retinas following DMSO and MTZ treatment. We found increased numbers of microglia in MTZ-treated eyes compared to DMSO controls at 48 hpt, but microglial numbers were not further impacted by the *w211* mutation (Figure 5 F). Morphology has historically been used as an approximation of microglial activation state (45, 46). Using such morphological features as an approximation of microglial state for this analysis, we defined “resting” microglia by features of visibly thin or elongated cell bodies and the presence of multiple extended cellular processes (Figure 5 G). We defined “activated” microglia as microglia cells with rounded and wider/thicker cell bodies with few-to-no processes extended (example image shown in Figure 5 G’). We also included a 3^rd^ category (collapsed microglia) for microglia with collapsed, compact, and mostly circular morphology (Figure 5 G’’), which may indicate apoptosis or other forms of cell death based on analysis from our previous work (47). We found that there were increased numbers of microglia with “activated” morphology following rod ablation (Figure 5 H), but the numbers of microglia with “activated” morphology were not significantly changed by the *nfe2l2a* loss-of-function mutation (Figure 5 H). Further, our morphological analysis did find significant numbers of collapsed microglia as a result of the rod ablation in *nfe2l2a*-sufficient or mutant retinas (Figure 5 I); while the p-value was statistically significant in MTZ-treated *w211+/-*eyes compared to DMSO controls, the mean number of collapsed microglia was approximately 2 per eye (Figure 5 I). We also found that microglia in MTZ-treated retinas, independent of *w211* genotype, had similar localization patterns with increased numbers of microglia localized to the outer nuclear/outer plexiform layers (Figure 5 J) where MTZ-ablated rods are located. Collectively, these results indicate that *nrf2a* mutant microglia may have increased ROS load but are still able to respond to rod ablation similarly to *nrf2a*-sufficient microglia.

We used RT-qPCR to assess the expression of selected genes that would be indicative of either redox imbalance, inflammation, endoplasmic reticulum (ER) stress, or glial reactivity following rod ablation at 48 hpt (Supplemental Figure 3). We examined expression of the *nfe2l2a* paralog *nfe2l2b*, which is a duplicate of *nfe2l2a* that has undergone subfunction partitioning (48) for potential compensatory expression with the loss of *nfe2l2a* function. Expression of *nfe2l2b* was not significantly increased in *w211-/-*mutants (Supplemental Figure 3). The gene sequestosome1 (*sqstm1/p62*) functions in autophagy and is associated with oxidative stress, acting both an activator of Nfe2l2 activity (via release from KEAP1) as well as a direct transcriptional target(49, 50). The expression of *p62* was significantly increased in MTZ-treated eyes with a functional copy of *nfe2l2a* but not in *nfe2l2a* loss-of-function mutants (Supplemental Figure 3A), likely indicating an effect of the loss of Nfe2l2a transcriptional activity. We did not detect changes in the expression of the antioxidant-related genes *nfe2l2a* and *gpx1a*, or in the ER stress response gene *bip* following MTZ treatment across both genotypes (Supplemental Figure 3). Further, the glial reactivity associated genes *gfap* and *vim* were strongly induced upon MTZ treatment or in *w211*-/-samples (Supplemental Figure 3). These RT-qPCR results further indicate that Nrf2 function is reduced in *w211*-/-retinas but suggest that the loss of *nfe2l2a* function alone does not result in similar alterations of redox biology comparable to what we previously found for microglia-deficient retinas (18).

### Loss of nrf2a function does not affect microglia or retinal cell death following the removal of MTZ

The loss of Nrf2 function has been reported to perpetuate the effects of damage and contribute to worsened outcomes following neural injury (51, 52), while Nrf2 overexpression has been shown to be protective of neuronal viability in degenerative conditions (53). In our previous study, we found increased numbers of TUNEL+ cells in multiple retinal layers of *rho:nfsb*-GFP; *irf8*-/-(microglia-deficient) retinas at 72 hpt (24 hours following MTZ-mediated rod ablation and washout) (18). To determine if loss of Nrf2 function resulted in similar impacts, we analyzed cell death via TUNEL staining in larval retinas following the same 24-hour recovery window after MTZ-mediated rod ablation in *w211+/-*and *w211-/-*mutants. We treated *rho*:nfsB-eGFP larvae with MTZ for 48 hours as previously, then removed them from MTZ treatment and collected samples 24 hours after MTZ washout (at 72 hpt). In addition to TUNEL staining, we also included immunofluorescent staining for microglia (Figure 6 A-D). The average number of TUNEL+ cells per retinal cryosection at 72 hpt was very few (Fig 6 H), with only occasional TUNEL+ cells localized to the ONL of MTZ treated samples (Fig 6 I), suggesting that the clearance of dying rods has occurred by this timepoint, consistent with our previous results for microglia-sufficient larval retinas (18). There was no impact of the *nfe2l2a*^*w211* genotype on TUNEL+ cell numbers in either DMSO or MTZ groups (Figure 6 H, I). The number of microglia per retinal section in MTZ-treated retinas remained modestly elevated compared to DMSO samples at this timepoint, but microglia numbers were not affected by the *w211* genotype (Figure 6 E). Further, we did not find differences between the two genotypes in the number of microglia with the previously described “activated” morphology (Figure 6 F), nor did we find significant differences in the number of microglia localized near the ONL/OPL (Figure 6 G). Since these samples included TUNEL staining, we scored collapsed, TUNEL+ microglia (4C4+ signal coupled with rounded, collapsed morphology, and positive for TUNEL signal, Fig 6 J). The numbers in all groups were minimal, with an average value in the elevated group of less than 1 cell per section (Figure 6 J). Collectively, these results indicate that the loss of *nfe2l2a* function did not result in prolonged impacts to microglia or retinal cell death.

**Figure 6:**
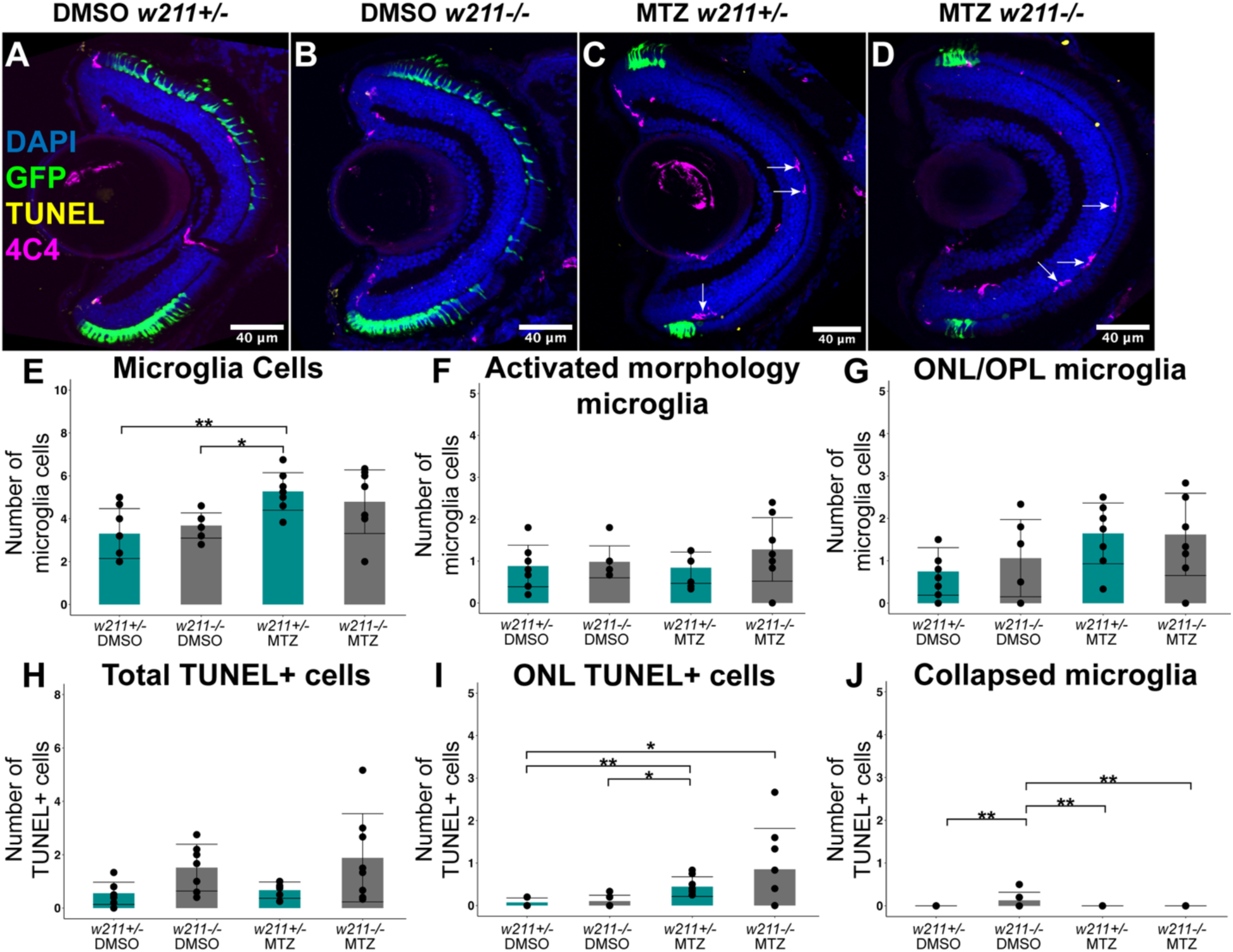
Retinal cell death and microglia analysis in *nrf2* loss-of-function mutants following MTZ washout. **A-D:** Representative images of retinal cryosections at 72 hpt showing DAPI (blue), GFP rods (green), TUNEL (yellow), and 4C4 (microglia stain, magenta) in DMSO (A, B) and MTZ-treated (C, D) larvae of either *w211+/-*(A, C) or *w211-/-*(B, D) genotypes. Arrows in C, D indicate microglia somata near the OPL/ONL. **E-G:** Average number of total microglia cells per retinal section (E), microglia with activated morphology (F), and ONL/OPL-localized microglia (G) per retinal cryosection (visualized with arrows in C-D). **H-I:** Average number of total TUNEL+ cells (H), ONL TUNEL+ cells(I), and TUNEL+, collapsed microglia (J) per retinal cryosection. Each datapoint represents the average number of cells per retinal cryosection of a single eye; 7-8 eyes per group (5-6 larvae). *p<0.05, **p<0.01, two-way ANOVA with Tukey’s HSD post-hoc test for multiple comparisons. For clarity, only significant differences are shown in these graphs.

### Nfe2l2a loss-of-function does not impact rod cell regeneration

Given that microglia are important for redox homeostasis following rod ablation (18), and microglia-deficient retinas show impacts to regenerated rods (Figures 2-4 above), we analyzed proliferation and rod regeneration in *nfe2l2a* mutant retinas (Figure 7 A-D’’’). These experiments again used *rho:nfsb*-eGFP siblings of either *w211*+/- or *w211*-/-genotype and similar stains to identify proliferating cells in retinal cryosections. Similar to the results described above, the proliferative response followed a similar trajectory and localization pattern, with overall PCNA+ numbers increasing by 96 hpt and most of the PCNA+ cells localized to the ONL rather than to Müller glia or in the INL (Figure 7E-H). A transitory difference between the MTZ-treated *w211+/-* and *w211-/-*groups was detected for PCNA+ cells in the INL and PCNA+ Müller glia at 72 hpt, where these numbers were reduced in *w211-/-*compared to *w211+/-*(Figure 7 G,H). There were no differences in numbers of PCNA+ cells in the ONL between *w211+/-* and *w211-/-*retinas at any timepoints following rod ablation (Figure 7 F). These results indicate that the loss of *nfe2l2a* function does not strongly impact proliferation of stem cell niches following rod ablation.

**Figure 7:**
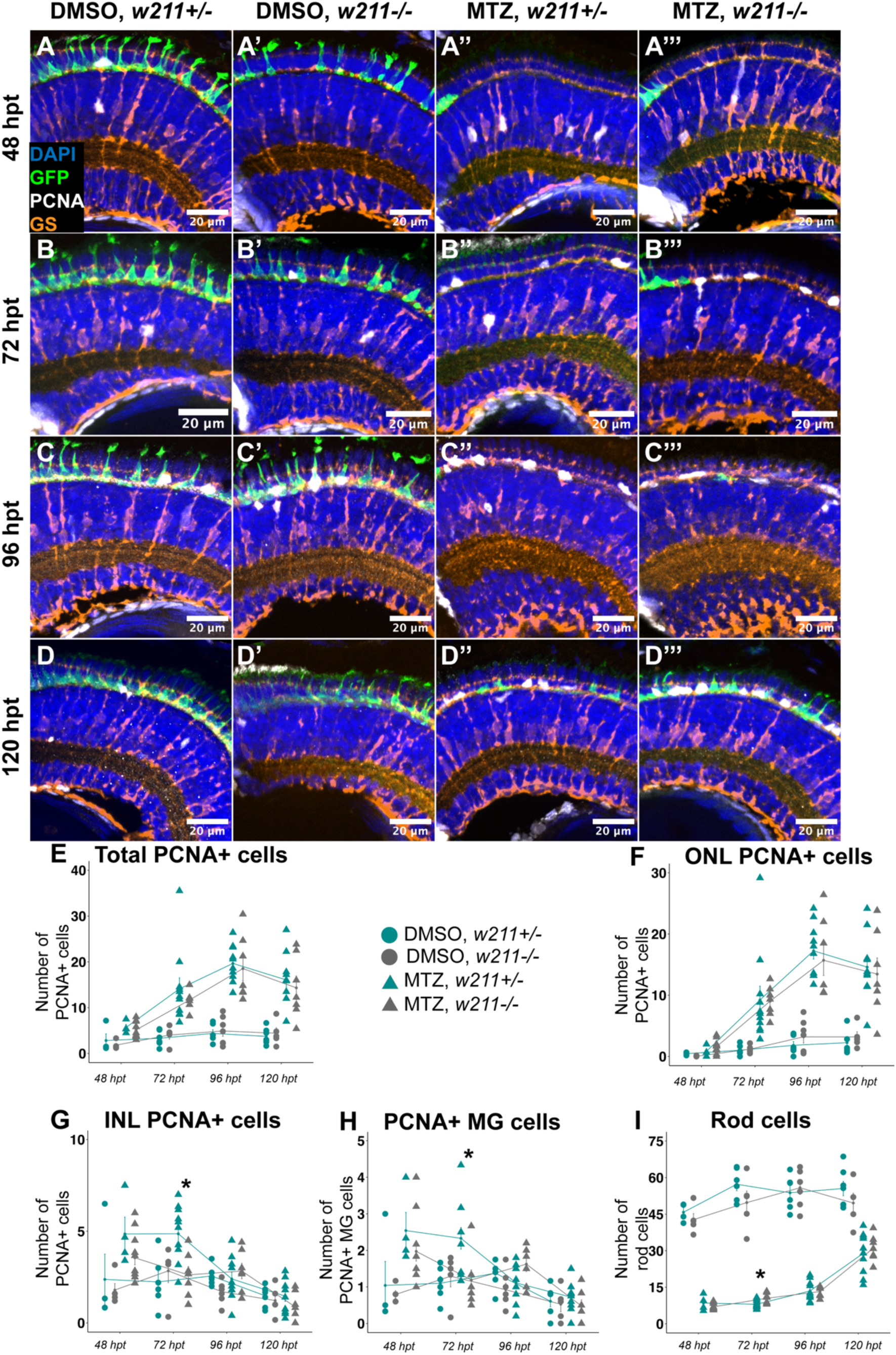
Cell proliferation following rod ablation in *nrf2* loss-of-function mutant retinas. **A-D’’’:** Representative images of dorsal region of retinas from retinal cryosections showing DAPI (blue), GFP+ rods (green), PCNA (white), and GS (orange) in DMSO (A-D, A’-D’) and MTZ-treated (A’’-D’’, A’’’-D’’’) and either *w211+/-*(A-D, A’’-D’’) or *w211-/-*(A’-D’, A’’’-D’’’). Top row shows samples at 48 hpt (A-A’’’); second row shows samples at 72 hpt (B-B’’’); third row shows samples at 96 hpt (C-C’’’), bottom row shows samples at 120 hpt (D-D’’’). **E-I:** Average number of total PCNA+ cells (E), ONL PCNA+ cells (F), and INL PCNA+ cells (G), PCNA+ Müller glia cells (H), and Rod cells per retinal cryosection over time from 48-120 hpt. Each point represents the average value of cells from retinal cryosections; 5-8 eyes; 3-5 larvae. Cyan circles denote DMSO *w211+/-*; gray circles denote DMSO *w211-/-*; cyan triangles denote MTZ *w211+/-*; gray triangles denote MTZ *w211-/-*. *Indicates significant difference between MTZ *w211+/-* and MTZ *w211-/-*where p<0.05, two-way ANOVA with Tukey’s HSD post-hoc test for multiple comparisons.

We performed analysis of regenerated rods in *w211+/-* and *w211-/-*retinas (Figure 8) using the same EdU immersion timing and detection method that was previously described for experiments with *irf8* larvae. Samples were collected at 216 hpt. Again, EdU+ GFP+ rods were not readily detected in DMSO-treated retinas (Figure 8 A,A’, B-C’, F-H) and EdU+ GFP+ rods were increased in MTZ-treated retinas (and primarily in dorsal region), but there were no differences in the number of EdU+ GFP+ rods between the *nfe2l2a* genotypes (Figure 8 A’’, A’’’, D-E’, F-H). Likewise, the only differences in total rod counts were a result of treatment and not *nfe2l2a* genotype, indicating that in contrast to the lack of microglia overall, the loss of *nfe2l2a* function did not impact the survival or numbers of regenerated rods (Figure 8 F-K).

**Figure 8:**
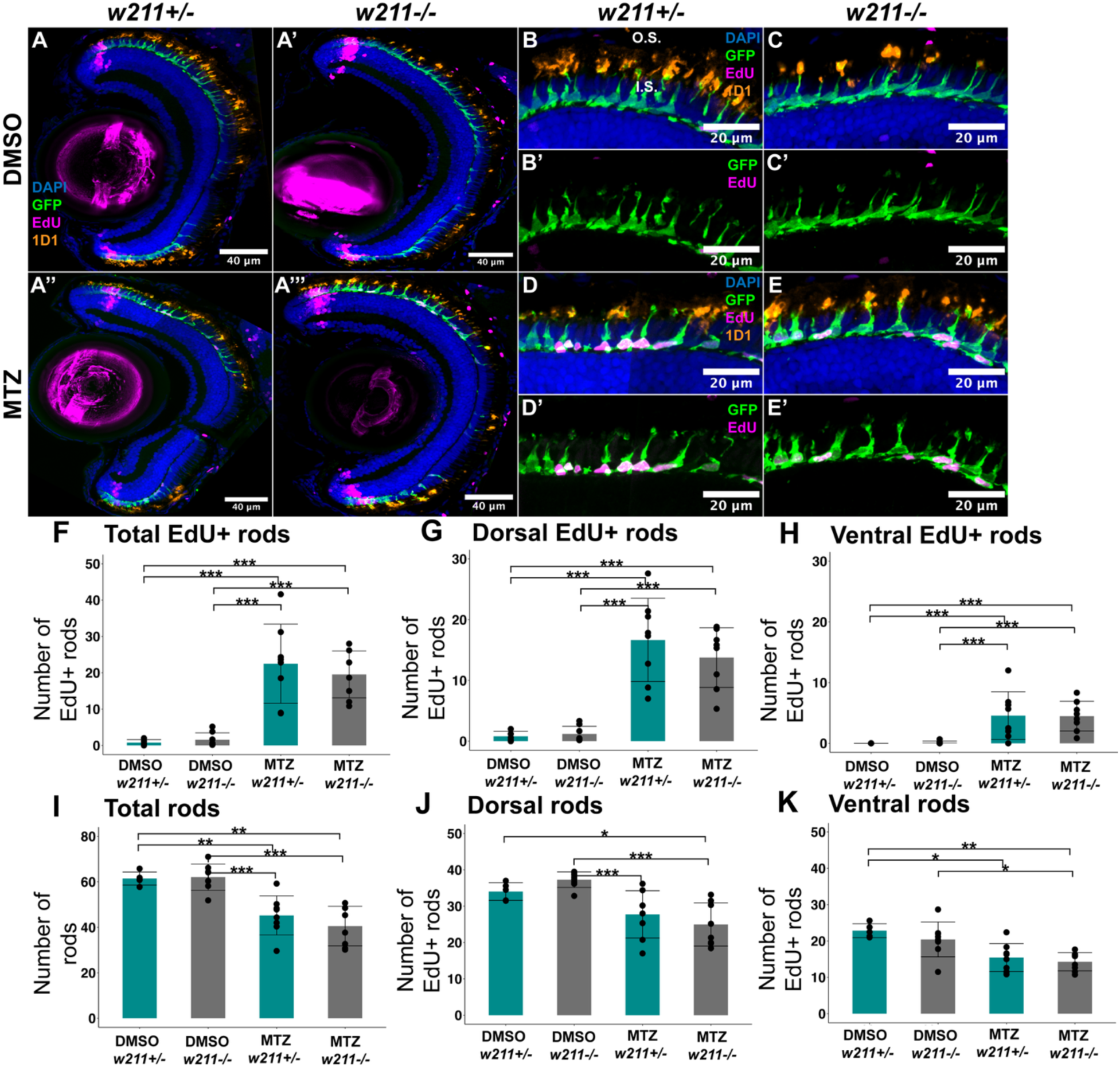
Loss of *nrf2* function does not affect numbers of regenerated rods. **A-A’’’:** Representative images of retinal cryosections at 216 hpt (9 days post-injury) from DMSO-(top row) and MTZ-treated retinas (bottom row) with either *w211+/-*(left-hand column) or *w211-/-*(right-hand column) genotype. Images show DAPI (blue), GFP rods (green), EdU (magenta), and 1D1 (rhodopsin, orange). **B-E’:** Dorsal regions of retinal cryosections at 216 hpt showing DAPI (blue), GFP rods (green), EdU (magenta), and 1D1 (rhodopsin, orange) (four-channel images B, C, D, E) or dual-channel images showing just GFP rods and EdU (B’, C’, D’, E’). Upper row shows DMSO-treated retinas with either *w211+/-*(B, B’) or *w211-/-*(C, C’) genotype. Lower row shows MTZ-treated (regenerating) retinas with either *w211+/-*(D, D’) or *w211-/-*(E, E’) genotype. O.S. and I.S. in B indicate outer segment and inner segment regions, respectively. **F-H:** Counts of EdU+ rods per retinal cryosection in the total retinal section (F), dorsal (G), and ventral (H) retina. Bars represent group means +/-SD, and each dot represents the average count per eye. **I-K:** Counts of total rods per retinal cryosection in the total retinal section (I), dorsal (J), and ventral (K) retina. Bars represent group means +/-SD, and each dot represents the average count per eye. n=5-10 eyes (3-6 larvae). *p<0.05, ** p<0.01, ***p<0.001, two-way ANOVA with Tukey’s HSD except Conover test with Bonferroni correction (H).

## Discussion

In this paper, we investigated the regeneration of rod photoreceptors in microglia-deficient *irf8*-/- and *nfe2l2a*-deficient larval zebrafish retinas. We found that the proliferation of progenitors following rod ablation is only modestly, and transiently, impacted in microglia-deficient retinas. However, microglia-deficient retinas contained reduced numbers of EdU+, regenerated rods at 216 hpt (9 days post-injury) and showed evidence of EdU+ rod photoreceptors that had failed to properly develop and instead had collapsed, abnormal morphology. The number of EdU+ rods at the earlier timepoint (144 hpt) was similar in both microglia-sufficient and -deficient retinas, but rods with abnormal, collapsed morphology were detected in *irf8* mutants at this earlier timepoint. Together, these results indicate that microglia support the proper development and/or survival of regenerated rod photoreceptors in the larval zebrafish retina. Given that total numbers of rods were similar in microglia-sufficient and -deficient retinas, the transient increase in proliferation seen at certain timepoints in *irf8-/-*retinas may represent an increase in progenitor production to mitigate the reduced number of surviving rods. In addition, since our experiments were performed in larvae, the overall number of rod photoreceptors may be influenced by the continual production of neurons during development (54), which could also be a factor in the overall rod numbers. In contrast to *irf8-/-*retinas, *nfe2l2a*-deficient retinas did not show significant impacts to proliferation after rod ablation or to rod regeneration. Together, these results indicate that Nrf2-independent functions of microglia likely sustain their redox function.

Our results are consistent with recent findings showing decreased numbers of EdU+ cells in microglia-deficient (*csf1ra -/-;csf1rb-/-*) retinas beginning at 10 days following retinal stab injury (15). Similar to our results, the decline in the number of EdU+ cells is not immediate but rather becomes evident at a later time. In another set of studies, retinal pigmented epithelium regeneration following NfsB-MTZ-induced ablation was found to be impaired in *irf8*-/- and *csf1ra/b*-/-mutant larval zebrafish (8, 55). In a previous larval rod ablation study (9), co-ablation of rods and microglia using the MTZ-NfsB system led to delayed regeneration of rods relative to the ablation of rods alone; however, recovery of rods over caught up to the rod-only ablation group at a later timepoint. This same study also found that immunosuppression by Dexamethasone prior to ablation treatment resulted in an impediment to regeneration and that the same treatment after the ablation treatment enhanced regeneration (9). These results suggest that microglial functions can have both positive and negative effects on regeneration and that an extant microglia population likely provides critical support early in the recovery process.

There were some different outcomes between the previous rod regeneration study (9) and our current study with regards to microglia effects that are worth consideration. The prior study (9) utilized inducible depletion and pharmacological immunosuppression to create a time-based study of microglia deficiency as opposed to our current system of genetic, developmental deficiency. There may be differences that could result from having a retina devoid of microglia-involved developmental processes as opposed to eliminating them at a later time. Recent work suggests a transcriptional reprogramming of the Muller glia when microglia are absent through development (15). Further, in White et al (9), YFP (rod) fluorescence volume of the whole larval retina was used to measure rod regeneration, whereas we used direct rod cell counts on retinal cryosections. We also included EdU labeling to identify regenerated rods following MTZ treatment during a selected time window. EdU labeling also allowed us to label the previous location of the CMZ at the time of EdU exposure. Because EdU exposure began at 48 hpt (when rod ablation has been executed and after which rod cell death subsides (18)), this allowed us to identify which regions of the retina contained regenerated rods from damage-induced progenitor niches. Together, this information allowed us to count rod cells most likely produced from regeneration as opposed to continuous development. We also found rods with an abnormal, collapsed morphology in *irf8-/-*retinas that were positive for both EdU and Rhodopsin. These were only found in regenerating regions and not CMZ-derived retina that developed after damage. Unfortunately, we were limited to analysis of fixed/static samples in our study because the larvae in our experiments were too large and pigmented for successful live timelapse imaging approaches; live imaging would provide necessary insight to the abnormal, collapsed rods detected in *irf8* mutants and could reveal whether the rods were generated then collapsed over time, or if they instead developed with abnormal morphology. Live imaging could also reveal if and when the abnormal rods are cleared from the retina.

Microglia may play other roles in facilitating the successful regeneration of rods in stages beyond the production of progenitors. Among these roles may be the remodeling of the extracellular matrix (ECM) and synapses, as microglia express genes associated with ECM remodeling following acute damage and during regeneration (56). ECM remodeling is an important element of neural development and is suggested to be important in regeneration as well, in part for the purposes of reorganizing the tissue for migration and integration of progenitors (57). The extent of Müller glia compensation towards ECM remodeling in these regards has yet to be determined, as do the full implications of microglia ECM remodeling, especially with regards to reintegration of rods into an already developed photoreceptor layer replete with cone cells in close, and highly organized, proximity. Microglia are also known to directly modify synapses in the CNS (58, 59) and can even influence neurotrophic factor production and resultant neuronal survival (60). All of these factors are not yet understood with regards to retinal regeneration and would need to be further investigated.

Though we did not detect obvious impacts to microglial morphology and viability, or to rod regeneration, in *nfe2l2a* loss-of-function mutants, this does not necessarily mean that antioxidant function is not important for regeneration. Indeed, there is redundancy with the antioxidant defenses that includes other activation mechanisms of antioxidant genes besides the activity of Nfe2l2a/Nrf2 (25, 61–64). We assessed the expression of the *nfe2l2a* paralog, *nfe2l2b,* and did not find increased *nfe2l2b* transcripts in *nfe2l2a* mutants, suggesting that this paralog does not provide significant compensatory function in the absence of *nfe2l2a*. This could be because Nfe2l2b is primarily involved with a different set of target genes and functions than Nfe2l2a (48, 65). However, other redox-responsive transcription factors could be activated, for example *nfe2l1a*, which encodes an ortholog of Nfe2l1/Nrf1. Nrf1 may have a degree of shared function and targets as Nfe2l2/Nrf2 and is also important for the antioxidant capacity of the cell (61, 62). Further, it has also been found that NF-kB can promote the transcription of antioxidant genes such as glutathione peroxidase 1 (Gpx1) and catalase (63). It is possible that these or other mechanisms of antioxidant activity could be activated to compensate for the loss of Nfe2l2a activity in microglia and in other retinal cell types. Although we detected greater intensity of CellRox™ in microglia following rod ablation in *nfe2l2a-*deficient retinas, we did not find changes in microglial numbers or the measured morphological characteristics. When considering the results from our previous study showing the significance of microglia redox function in the retina (18), and the impact to regenerated rod survival when microglia are absent (this paper), our results likely indicate that microglia are robust in their use of mechanisms towards redox homeostasis. Further, the robustness of microglia antioxidant defenses, including the glutathione-dependent enzymes detected in our previous studies (6, 18), may provide insight as to why attempts to directly ablate NfsB-expressing microglia using MTZ are often challenging (47, 66).

Another important consideration with our experimental outcomes and the *rho*:nfsb-eGFP transgenic line for MTZ-mediated ablation of rod photoreceptors is the biology of rod photoreceptor replacement in zebrafish, which can come from MGPCs and rod precursor cells (30, 39). The initial experiments with the *rho*:nfsb-eGFP^nt19 line used adult fish and elicited significant levels of Müller glial proliferation owing to extensive rod ablation (30). In larval retinas, we did not detect abundant Müller glial cell proliferation following rod ablation even though rod cells were extensively ablated. Instead, cell proliferation was more abundant in the ONL, suggesting that rod precursors supply the predominant proliferative response (39). MTZ treatment of *gnat2*:nfsb-mCherry transgenic larval fish (67), in which highly abundant *gnat2*-expressing cone photoreceptors are ablated upon MTZ exposure, elicited proliferation of Müller glia and MGPC (67, 68). This is not to discard the value of this rod ablation system, but the observed lack of a strong Müller glia response seen following rod ablation in larvae is a key element to consider when interpreting the results of our study and to consider microglial support of regeneration. Further, in our recent paper involving regeneration experiments using *irf8* mutants and *gnat2*:nfsb-mCherry larval fish, we did not detect impacts to the regeneration of cones in *irf8* mutants (41) as we did here in this work for rods. We reconcile these different outcomes by considering the following. First, upon ablation of cones in larval *irf8* mutant retinas, we detected compensatory immune cell populations that responded to cone death (41), while microglia remained essentially absent from *rho*:nfsb-eGFP, *irf8* mutant retinas following rod ablation, with a persistence of residual TUNEL+ cells indicative of professional phagocyte-deficiency (18). The compensatory leukocytes responding to cone damage could provide functions that support cone survival, including redox functions natively provided by microglia, and these compensatory cell populations may be more strongly evoked to contribute to the restoration of redox balance. In addition, distinct stem cell niches, and even photoreceptor types themselves, may be more or less sensitive to redox imbalance.

While the presence of microglia may not be absolutely essential for regeneration to proceed following retinal damage (11), this may be because of compensatory functions or responses from other cells (11, 41, 69), which have yet to be clearly defined in multiple microglia-deficient contexts (including regeneration). Our work here suggests that additional rods may be produced to replace those that do not survive in *irf8* mutant retinas, resulting in a similar level of rod photoreceptors overall. This could be part of the regenerative response and/or an adaptation of development (70). Importantly, lack of essentiality does not ultimately contradict reports of microglia function in and impact on retinal regeneration (8, 15, 17, 71). While individual molecular components may not be a “make-or-break” for regeneration outcomes, the initiation of MG-derived regeneration and the production of surviving regenerated neurons may be influenced by multiple molecular mechanisms (15). Altogether, our results reflect the principle of redundancy in cell biology, which is ultimately a vital component of survival given that cellular and tissue components are bound to incur damage in environments where they are regularly challenged by ROS and other potential sources of damage.

## Materials and Methods

### Animals

All procedures were conducted with protocols approved by the Institutional Animal Care and Use Committee at the University of Idaho. Adult zebrafish (*Danio rerio*) were housed in an aquatic vivarium with filtered, recirculating water, maintained at 28.5°C, and an automated 14h light:10 h dark cycle following procedures outlined in *The Zebrafish Book* (72). Zebrafish embryos were collected into glass beakers from off-system spawn tanks following morning light onset, which is considered the approximate time of fertilization (0 hours post-fertilization). Because zebrafish cannot be sexed before reproductive maturity, sex of larval zebrafish used in the experiments could not be determined and was therefore not factored. Embryos that were used for whole eye imaging were housed in a Heratherm™ incubator, with 14:10h light:dark cycle, at 28°C and treated with N-Phenylthiourea (PTU) at a final concentration of 0.006% in order to prevent the development of pigmentation. The water and PTU were refreshed each day. Prior to collection of larvae at experimental endpoints, larvae were terminally anesthetized in a working solution of 0.25 mg/mL MS-222/tricaine (Syndel USA).

### Zebrafish lines

The transgenic line *rho*:nfsb-eGFP*^nt19* was generated previously and gifted by Dr. David Hyde’s lab (University of Notre Dame) (30). In this line the *rho* (rhodopsin) promoter drives the expression of nfsB fused to eGFP fluorescent protein in rod photoreceptors. Carriers of the *nt19* transgene were identified by GFP signal visible in eyes at age 3 days post fertilization (dpf) when viewed under a fluorescent microscope. The *nt19* line was outcrossed to our wild-type strain referred to as SciH, originally obtained from Scientific Hatcheries (now Aquatica Tropicals), and to *irf8*^st95 and *nfe2l2a^w211* carriers. The *irf8^st95* line was generated and gifted previously by Dr. Celia Shiau (University of North Carolina) (34). Genetic crosses were performed to produce fish carrying the *rho*:nfsB-eGFP transgene and heterozygous or homozygous for the *irf8*^*st95* allele; these are referred to throughout the manuscript as *irf8+/-* and *irf8-/-*, respectively. The *nfe2l2a*^w211 line was generated previously by the Gallagher Lab (University of Washington) (35) and obtained from the Zebrafish International Resource Center (ZIRC). Similar to the *irf8* crosses, the *w211* allele was crossed to produce *rho:*nfsB-eGFP+ fish either heterozygous or homozygous for the *w211* allele, referred to as *w211+/-* or *w211-/-*, respectively. Experiments were performed with clutches of both *irf8* and *w211* genotypes, and genotype unknown (blinding) at the time of the experimental procedures. Genotype of the larvae was determined post-fixation from tail clips using a PCR/Restriction digest assay as described in (34) (*irf8*), and *w211* genotyping was performed using primers previously reported (35) (FP: 5’-AGGAGCAGGAGAAGACACTG-3’; RP: 5’-GGCATTTGTGTCAAATCTCACAGG-3’, annealing temperature 63° C); however, we also used an HhaI restriction enzyme digest (1 hour, 37° C) of the PCR product, as the HhaI cut-site is missing from the mutant allele.

### Metronidazole treatment

For rod ablation, treatment of *rho*:nfsb-eGFP larval zebrafish began at the age of 3.5 dpf. Larvae were immersed in a solution of 10mM metronidazole (MTZ) in 0.1% by volume dimethyl sulfoxide (DMSO), or in a solution of 0.1% DMSO (vehicle), prepared in fish system water. Solutions were prepared at least four hours prior to use, followed by constant mixing, and were protected from light. Treatments were performed in a Heratherm™ incubator set at 28°C. Due to the light-sensitivity of MTZ, larvae exposed to DMSO/MTZ solutions were treated in petri dishes and protected from light by covering dishes with aluminum foil. After 24 hours of treatment, larvae received a fresh solution of their respective treatment and then returned to the incubator for another 24 hours. Following the duration of treatment (48 hours post treatment, hpt), larvae were removed from dishes and either fixed for tissue preparations (described below) or transferred to a pulse of CellRox™ immersion. For samples collected at timepoints after 48 hpt, larvae were transferred to clean water for recovery and collected at their designated experimental endpoint with water changes and feeding.

### CellRox™ immersion

At the end of the DMSO/MTZ treatment duration, larvae were removed from their respective treatment solutions and placed in a solution consisting of 0.1% CellRox™ Deep Red reagent (Invitrogen, working concentration of 2.5 µM) and 0.006% N-phenylthiourea in fish system water. Larvae were incubated in CellRox™ solutions for 30 minutes at 28°C and protected from light. Following incubation, larvae were removed from solution, terminally anesthetized in a working solution of 0.024% MS222-tricaine, then fixed in a solution of 4% paraformaldehyde in 1x PBS overnight at 4°C. Following fixation, samples were processed according to the whole-tissue immunofluorescence staining protocol as described below and reported in our previous work (18).

### Immunofluorescence of whole tissue

At designated experimental endpoints, larvae were fixed in a solution of 4% paraformaldehyde in 1x phosphate buffered saline (PBS) overnight at 4°C. The next day, the fixation solution was removed, and larvae were rinsed once in 1X PBS+ 1% Triton-X100 (PBST) then three times in 1X PBS for 5 minutes each. Following PBS rinses, the larvae were dehydrated with four 100% methanol washes for 10 minutes each, then stored in 100% at −20°C. Prior to staining, methanol was removed from the samples, and they were rehydrated with two 15-minute washes of 1x PBST (1% triton-X-100). Following rehydration, the blocking step was performed by incubation in antibody dilution buffer (1% normal donkey serum, 0.1% of sodium azide, 0.1% triton-X-100 in 1X PBS) for at least one hour at room temperature. The primary antibody solution was prepared in antibody dilution buffer. The primary antibody used was mouse 4C4 (1:100, hybridoma supernatant from clone 7.4.C4). Samples with primary antibody mixtures were incubated overnight at 4°C with constant rocking. Following primary antibody incubation, the samples were washed with 500 µL of 1x PBST (1% triton-X-100) for 30 minutes at room temperature. Secondary antibody solutions were prepared at 1:200 dilution in antibody dilution buffer. The secondary antibodies used in these experiments was goat anti-mouse Cy7 (1:200, AAT Bioquest). DAPI was included in the secondary antibody solution at a dilution of 1:1000. Samples were incubated in secondary antibody solution overnight at 4°C with constant rocking. The next day, the samples were washed again with 500µL of 1x PBST for 30 minutes at room temperature, followed by a 20-minute rinse with 1x PBS.

For imaging, eyes were dissected from fixed and stained whole larvae and suspended in glycerol. A “coverslip sandwich” was made using a 22×60 mm microscope slide coverslip and attaching two smaller 22.5×22.5 mm coverslips on each end of the longer coverslip using adhesive. Samples were mounted in the space on the 22×60 mm coverslip between the 22.5×22.5 mm coverslips. When the samples were oriented, a second 22×60 mm coverslip was attached by placing on top of the smaller coverslips to create a “sandwich” effect.

### Preparation of retinal cryosections

At designated experimental endpoints, larvae were fixed in solution of 4% paraformaldehyde in a 5% sucrose phosphate buffer and incubated overnight at 4°C. Afterwards, the fixative was removed, and the samples were washed in 5% sucrose phosphate buffer thrice for 10 minutes each. The samples were then washed with graded series of sucrose phosphate solutions beginning with a 2:1 ratio of 5%:20% sucrose phosphate buffer, followed by 1:1, and 1:2 ratio for 30 minutes each. Finally, the samples were incubated with a 20% sucrose phosphate buffer overnight at 4°C. Samples were then embedded into tissue blocks composed of a medium of 2:1 20% sucrose phosphate buffer: OCT and stored at −20°C at least overnight. Tissue blocks were sectioned using a Leica CM3050 cryostat with 10µm cryosection thickness, dried in a desiccator overnight, then stored at −20°C.

### TUNEL assay and immunostaining of retinal cryosections

Microscope slides with retinal cryosections were thawed for 10 minutes at room temperature following removal from the freezer. Because of the extant GFP fluorophore present in the tissue, the samples were protected from light as much as possible during active procedures and during all incubation steps. Detection of PCNA requires an antigen retrieval step, so we performed a previously described antigen retrieval procedure using a heated citrate buffer for 20 minutes (73). Following antigen retrieval, the immunostaining procedure was carried out as previously described, but with different antibodies. For primary antibodies we used rabbit anti-PCNA (Proteintech 10205, 1:200) to detect PCNA and mouse anti-glutamine synthetase (BD Biosciences 610517, 1:200) to detect Müller glia. Because the antigen retrieval process damages the transgenic fluorophore, we used chicken anti-GFP (Abcam ab13970, 1:1000) to recover visualization of rod photoreceptors. For secondary antibodies we used anti-rabbit Alexa Fluor™ 647 (Jackson Immunoresearch, 1:200), goat anti-mouse Cy7 (AAT Bioquest, 1:200), donkey anti-chicken Alexa Fluor™ 488 (Jackson Immunoresearch, 1:200), as well as DAPI (1:1000) to visualize cell nuclei.

Tissue was blocked for at least 1 hour at room temperature in blocking solution (described above), then incubated with primary antibody solution at 4°C overnight. Primary antibody solution was prepared in antibody dilution buffer with mouse anti-4C4 (1:100, hybridoma supernatant from clone 7.4.C4). The next day, slides were washed in 1x PBST (1% Triton-X-100) for 30 minutes at room temperature. The secondary antibody solution was prepared with donkey anti-mouse Alexa Fluor™ 647 (Jackson Immunoresearch, 1:200) and DAPI at a dilution of 1:1000. Following the PBST wash, the slides were incubated with secondary antibody solution for 2 hours at room temperature. Slides were once again washed with 1x PBST for 30 minutes at room temperature, followed by a 20-minute wash in 1x PBS. Stained tissue was mounted in Vectashield Vibrance® (Vector Labs), covered with microscope slide coverslips, and cured = in the dark at room temperature for at least two hours prior to imaging.

### EdU exposure and detection

Preparation of stock solutions: 5-ethynyl 2-deoxyuridine (EdU) was acquired through the Invitrogen Click-It EdU kit with Alexa Fluor 647 (C10340) prepared as a 10 mM stock solution in DMSO and stored at −20°C until needed. For EdU treatment, a final working solution of 125 µM EdU in fish system water was prepared from the stock solution, and larvae were immersed in solution for 24 hours from 48-72 hpt (5.5-6.5 dpf). Following treatment, larvae were rinsed with system water to remove EdU solution and were set to continue recovery with daily water changes and the administration of larval food. Larvae were collected at either 144- or 216 hpt according to the fixation protocol for cryosections. Samples were processed according to the previously described cryosection preparation procedure.

Components of the Click-It EdU kit were prepared according to manufacturer’s directions, including 10X reaction buffer (stored at 4C), 10x reaction additive (stored at −20°C in aliquots). Because copper sulfate and the Alexa Fluor 647 azide reagents were consumed faster than the other reagents, we acquired additional reagents according to recommended products from the manufacturer prior to our experiments. Additional copper sulfate was purchased (Sigma Aldrich, C1297) and prepared as 100 mM stock solution in water. Further, additional Alexa Fluor 647 azide was purchased as a triethylammonium salt (Invitrogen, A10277). A stock solution of 1 mM was prepared by adding 588 µL of DMSO to the salt. Because the final concentration of Alexa Fluor 647 azide provided in the Click-It kit is not provided, we used the separately prepared Alexa Fluor 647 azide for all of our experiments to ensure equal concentration across all samples. For detection of EdU, slides were thawed at room temperature for 10 minutes. Due to the presence of GFP fluorophore, slides were protected from light exposure in this procedure. Slides were individually covered with 1mL 1X PBS for 10 minutes. Then, a liquid blocker gel was applied to the outline of the slides, and the slides were covered in 1X PBST (0.5% Triton-X-100) for 20 minutes at room temperature, then rinsed in 1X PBS. The EdU detection mixture was prepared with the following portions of reagent per each slide: 430µL of 1X reaction buffer (from 10X stock), 20 µL of copper sulfate (from the 100 mM stock), 50 µL 1X reaction buffer additive (from 10x stock), and 1.25 µL Alexa Fluor 647 azide (from 1 mM stock). Slides were then transferred to a humidified slide container and were overlayed with 500 µL of the EdU detection mixture each and left to incubate at room temperature for 30 minutes. Following incubation with the detection mixture, slides were rinsed thrice with 1X PBS. From this point, slides would then follow the immunofluorescence procedure for cryosections previously described.

### Confocal Microscopy

Images were acquired on a Nikon Eclipse Ti2-E, Crest X-light V3 spinning disk confocal microscope using a CFI APO LWD 40x water immersion 1.15 NA λ S DIC N2 objective. Image acquisition was performed using a computer workstation running Nikon Elements software. For whole eyes, image stacks were acquired with 3μm z-steps. For retinal cryosections, images were acquired with 2µm z-steps. DAPI was imaged using the DAPI laser line (365 nm). GFP and Alexa Fluor™ 488 were imaged using a GFP laser line (∼480 nm). TUNEL TMR Red was imaged using orange/red laser line (∼560 nm). CellRox™ Deep Red dye and Alexa Fluor™ 647 were imaged done using excitation with a far-red laser (∼640 nm). Imaging of the Cyanine 7 (Cy7) fluorophore was done with a Cy7 excitation laser line (∼711nm).

### Image viewing and preparation

Image processing and quantifications were performed using ImageJ/FIJI. Viewing of both individual z-stacks and maximum projection images of select z-stacks was used to analyze according to the intended metric. A cell counter plug-in was used to count cells of interest (e.g., TUNEL+ cells). Microscopy images presented in our figures were converted into RGB tiff files from their original 16-bit format.

### Quantification of fluorescence intensity

Quantification of fluorescence intensity in confocal images was performed using FIJI/Image J. Fluorescence intensity was determined using single z-stack images and was adapted from our lab’s previous publication (69) and used previously (18). Prior to measurement, the measurement options in Image J were set to include area and integrated density. For CellRox™ fluorescence, the intensity of the CellRox™ signal was measured by drawing a region of interest (ROI) around a microglial cell determined by the shape of the signal of the 4C4 antibody stain. For the intensity of mRNA transcript via HCR-FISH, the bounds of the microglia were approximated by the bounds of the *mpeg1* transcript signal. The measured intensity of the target signal of interest was then measured within those bounds to serve as the approximate intensity within microglia. The approximate median z-stack of a cell was chosen for fluorescence measurement. In addition to the fluorescence intensity within the bounds of the cell, several regions of background outside the cell were measured and averaged to provide mean background signal value. From here, the average background was multiplied to the area of the region of interest to generate a value of mean background signal. That generated value was subtracted from the integrated density to yield the corrected total cell fluorescence (CTCF).

### Categorical scoring of microglia in confocal images

Categorical classification of microglia was performed by examining all z-stack confocal images acquired to determine the shape and features of each microglial cell. This was done for both whole eye images and images of retinal cryosections. Microglia were scored into morphophysiological categories. “Resting” microglia were defined by features of visibly thin or elongated cell bodies and the presence of multiple extended cellular processes (see Figure 5 G). “Activated” microglia were defined as microglia cells with rounded and wider/thicker cell bodies with few-to-no processes extended (see Figure 5 G’). “Collapsed microglia” were defined as microglia with collapsed, compact, and mostly circular morphology (see Figure 5 G’’). In a subset of analyses, TUNEL labeling was also included for “collapsed microglia”. For the ONL-localization score of microglia, we identified the position of the cell body relative to DAPI staining of the ONL. This category included cells that were in the ONL/photoreceptor layer or had cell bodies positioned perpendicular to the ONL and were approximately 5 or fewer micrometers basal to the ONL and/or had visible process extension into the ONL (See arrows in Figure 6 for example).

### RT-qPCR

At the experimental endpoint, anaesthetized larvae were placed in a 1.5mL Eppendorf tube, water was removed, and larvae were flash frozen on liquid nitrogen and immediately overlayed with super-chilled 100% methanol. Samples were stored in a −80°C freezer in methanol until ready for RNA extraction. For eye dissection, the samples were held on ice and eyes were removed from larval heads and placed into RNA lysis buffer (Zymo Quick-RNA microprep kit). Tissue was homogenized in lysis buffer using a handheld homogenizer (Bio-Gen Series PRO200, ProScientific). RNA was extracted Zymo Quick-RNA microprep kit (Zymo Research) according to the manufacturer’s protocol. Each sample was a pool of both eyes from a single fish. For genotyping of samples, the remainder of the body was transferred to 25µl NaOH and gDNA prepared as established in *Meeker et al.*(74). DNA was used for genotyping as described above. Following RNA extraction, cDNA was prepared using SuperScript IV cDNA synthesis kit (Invitrogen). cDNA was stored at −20°C until use.

Quantitative PCR (qPCR) was performed using PowerTrack SYBR Green Master Mix (Applied Biosystems), with 1 ng of cDNA per reaction. Reactions were performed in duplicates. Primers for genes of interest were selected from previous publications and are listed in **Table 1**. The qPCR reaction was carried out in a QuantStudio™ 3 Real-Time PCR System (ThermoFisher) under standard reaction program settings with 40 amplification cycles. *β*-actin served as the housekeeping gene for our analysis, and fold-change was determined by the ΔΔCT method.

**Table 1:**
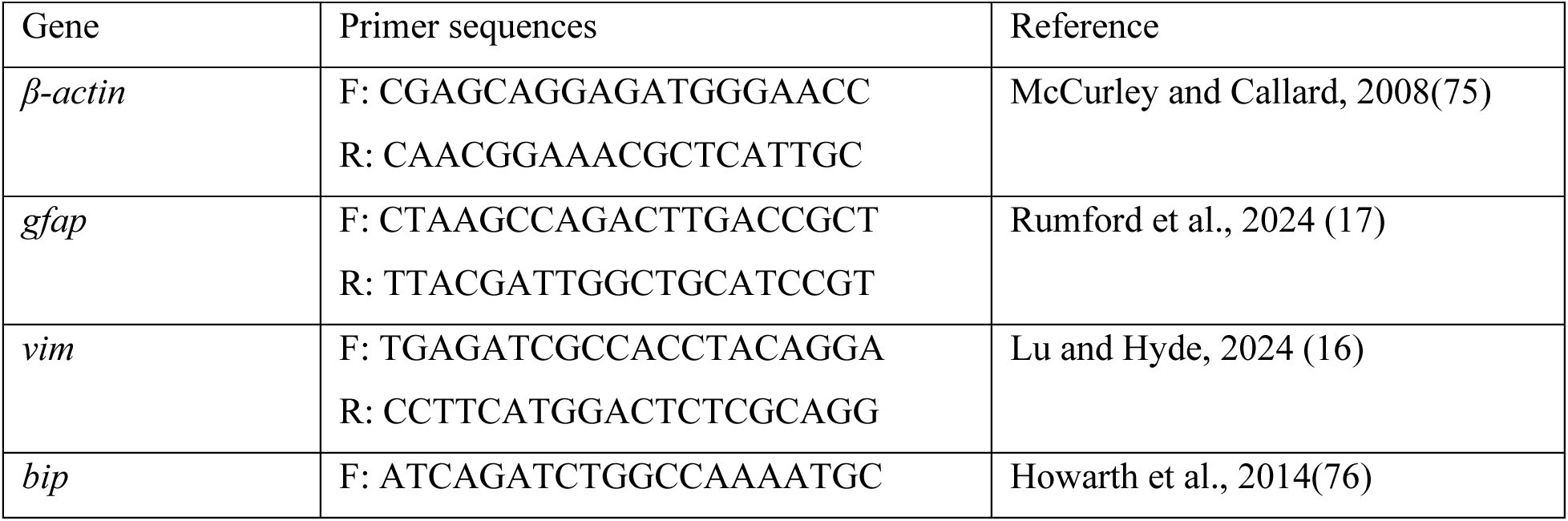
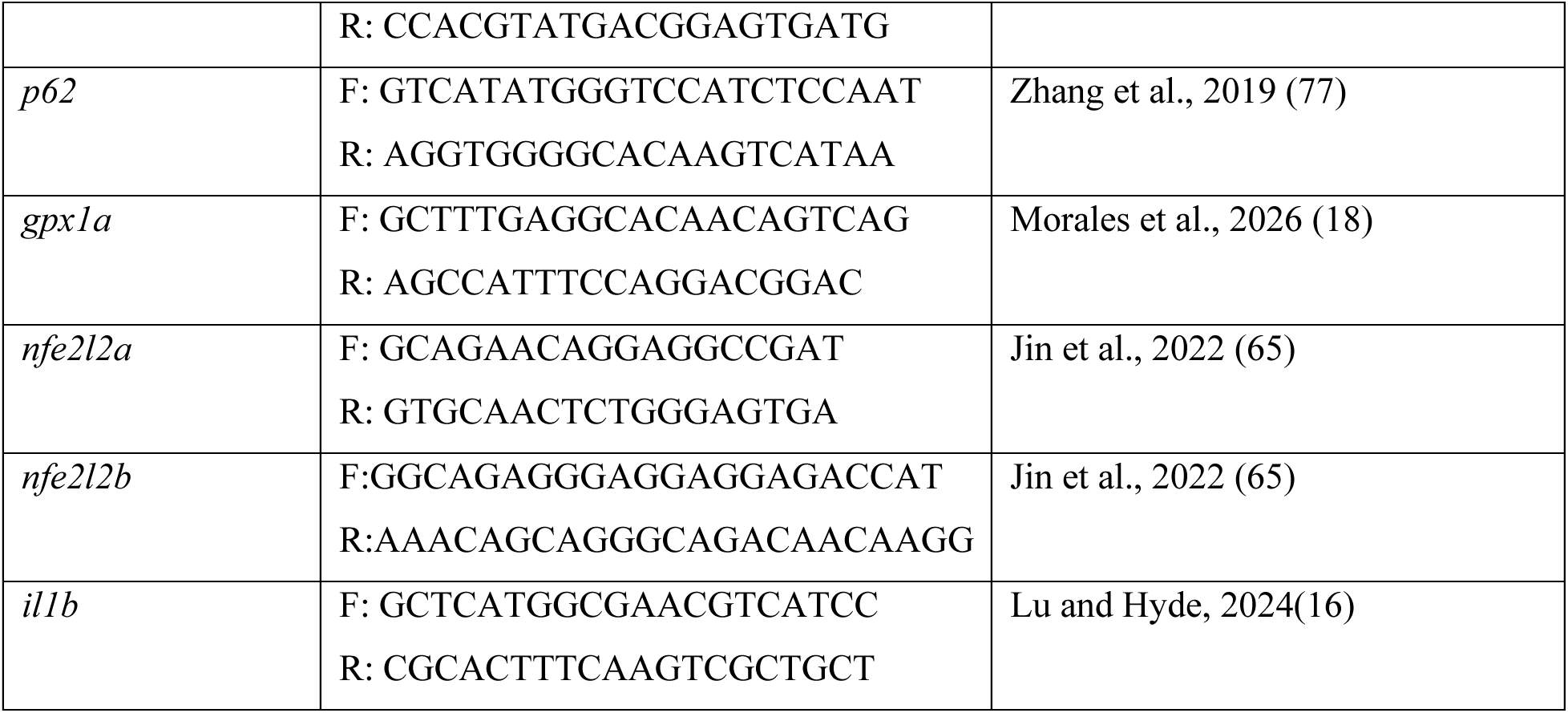
Primers used for qPCR.

### Statistical Analysis

Statistics were computed in R-Studio. For comparisons between multiple groups, Shapiro-Wilk test was used to check data for normality (parametric distribution). For non-normal data, the “BestNormalize” package for R was used to transform a dependent variable before performing statistical analysis. The transformed data was once again tested for normality using Shapiro-Wilk test. A two-way ANOVA test was then performed on successfully transformed, normalized data, followed by Tukey’s honest significant difference post-hoc test. P-values less than 0.05 were considered statistically significant. For data that could not be normalized with our transformations, we performed a Conover’s Test with Bonferroni correction, and p-values <0.025 were considered significant (α/2). Actual p-values are reported for all statistical tests in Supplemental Information.

## Acknowledgments

We thank Dr. David Hyde for gifting the *rho:nfsB-eGFP^nt19* transgenic zebrafish, as well as Dr. Celia Shiau for gifting the *irf8^st95* mutant zebrafish. We thank the University of Idaho Institute of Modeling Complex Interactions(IMCI) Image and Data Acquisition Core (IDAC),, and Institute for Interdisciplinary Data Science for equipment and logistical support. We thank Parker Van Dam (University of Idaho) for providing technical support and Jordan Rumford for care of zebrafish. We thank Dr. Deborah Stenkamp (University of Idaho) for providing critical review of an earlier version of the manuscript.

## Author contributions

Manuscript drafting and editing: M.M., D.M.M.; visualization: M.M., D.M.M.; investigation: M.M., D.M.M.; formal analysis: M.M.; conceptualization: M.M., D.M.M.; supervision: D.M.M.; project administration: D.M.M.; funding acquisition: D.M.M.

## Bibliography

1. Fausett, B. V., and Goldman, D. (2006) A Role for 1 Tubulin-Expressing Muller Glia in Regeneration of the Injured Zebrafish Retina. Journal of Neuroscience. 26, 6303–6313

2. Bernardos, R. L., Barthel, L. K., Meyers, J. R., and Raymond, P. A. (2007) Late-Stage Neuronal Progenitors in the Retina Are Radial Muller Glia That Function as Retinal Stem Cells. Journal of Neuroscience. 27, 7028–7040

3. Thummel, R., Kassen, S. C., Enright, J. M., Nelson, C. M., Montgomery, J. E., and Hyde, D. R. (2008) Characterization of Müller glia and neuronal progenitors during adult zebrafish retinal regeneration. Exp. Eye Res. 87, 433–444

4. Fimbel, S. M., Montgomery, J. E., Burket, C. T., and Hyde, D. R. (2007) Regeneration of Inner Retinal Neurons after Intravitreal Injection of Ouabain in Zebrafish. The Journal of Neuroscience. 27, 1712–1724

5. Nagashima, M., Barthel, L. K., and Raymond, P. A. (2013) A self-renewing division of zebrafish Müller glial cells generates neuronal progenitors that require N-cadherin to regenerate retinal neurons. Development. 140, 4510–4521

6. Mitchell, D. M., Sun, C., Hunter, S. S., New, D. D., and Stenkamp, D. L. (2019) Regeneration associated transcriptional signature of retinal microglia and macrophages. Sci. Rep. 10.1038/s41598-019-41298-8

7. Todd, L., Finkbeiner, C., Wong, C. K., Hooper, M. J., and Reh, T. A. (2020) Microglia Suppress Ascl1-Induced Retinal Regeneration in Mice. Cell Rep. 10.1016/j.celrep.2020.108507

8. Leach, L. L., Hanovice, N. J., George, S. M., Gabriel, A. E., and Gross, J. M. (2021) The immune response is a critical regulator of zebrafish retinal pigment epithelium regeneration. PNAS. 118, e2017198118

9. White, D. T., Sengupta, S., Saxena, M. T., Xu, Q., Hanes, J., Ding, D., Ji, H., and Mumm, J. S. (2017) Immunomodulation-accelerated neuronal regeneration following selective rod photoreceptor cell ablation in the zebrafish retina. Proc. Natl. Acad. Sci. U. S. A. 114, E3719–E3728

10. Mitchell, D. M., Lovel, A. G., and Stenkamp, D. L. (2018) Dynamic changes in microglial and macrophage characteristics during degeneration and regeneration of the zebrafish retina. J. Neuroinflammation. 15, 163

11. Song, P., Parsana, D., Singh, R., Pollock, L. M., Anand-Apte, B., and Perkins, B. D. (2024) Photoreceptor regeneration occurs normally in microglia-deficient irf8 mutant zebrafish following acute retinal damage. Sci. Rep. 10.1038/s41598-024-70859-9

12. Xu, J., Li, Y., Li, X., Tan, X., Liu, L., Cao, L., and Xu, H. (2025) Microglia-Derived IL-6 Promotes Müller Glia Reprogramming and Proliferation in Zebrafish Retina Regeneration. Invest. Ophthalmol. Vis. Sci. 10.1167/iovs.66.4.67

13. Palazzo, I., Todd, L. J., Hoang, T. V., Reh, T. A., Blackshaw, S., and Fischer, A. J. (2022) NFkB-signaling promotes glial reactivity and suppresses Müller glia-mediated neuron regeneration in the mammalian retina. Glia. 70, 1380–1401

14. Emmerich, K., White, D. T., Kambhampati, S. P., Casado, G. L., Fu, T. M., Chunawala, Z., Sahoo, A., Nimmagadda, S., Krishnan, N., Saxena, M. T., Walker, S. L., Betzig, E., Kannan, R. M., and Mumm, J. S. (2023) Nanoparticle-based targeting of microglia improves the neural regeneration enhancing effects of immunosuppression in the zebrafish retina. *Commun*. Biol. 10.1038/s42003-023-04898-9

15. Mitra, S., Devi, S., Lee, M. S., Sahu, A., Jui, J., Kriukov, E., Baranov, P., and Goldman, D. (2026) Müller glia-microglia cross talk reprograms the Müller glia transcriptome for cell division-related processes during retina regeneration. Proc. Natl. Acad. Sci. U. S. A. 10.1073/pnas.2535044123

16. Lu, C., and Hyde, D. R. (2024) Cytokines IL-1β and IL-10 are required for Müller glia proliferation following light damage in the adult zebrafish retina. Front. Cell Dev. Biol. 10.3389/fcell.2024.1406330

17. Rumford, J. E., Grieshaber, A., Lewiston, S., Reed, J. L., Long, S. S., and Mitchell, D. M. (2024) Forced MyD88 signaling in microglia impacts the production and survival of regenerated retinal neurons. Front. Cell Dev. Biol. 10.3389/fcell.2024.1495586

18. Morales, M., and Mitchell, D. M. (2026) Microglia maintain retinal redox homeostasis following ablation of rod photoreceptors in zebrafish. Journal of Biological Chemistry. 10.1016/j.jbc.2026.113169

19. Weimar, H. V., Farre, A. A., Rumford, J. E., and Mitchell, D. M. (2026) Müller glial and microglial responses coupled to recovery of cone photoreceptors following limited cone ablation in zebrafish retina. Sci. Rep. 10.1038/s41598-026-40141-1

20. De Almeida, A. J. P. O., De Oliveira, J. C. P. L., Da Silva Pontes, L. V., De Souza Júnior, J. F., Gonçalves, T. A. F., Dantas, S. H., De Almeida Feitosa, M. S., Silva, A. O., and De Medeiros, I. A. (2022) ROS: Basic Concepts, Sources, Cellular Signaling, and its Implications in Aging Pathways. Oxid. Med. Cell. Longev. 10.1155/2022/1225578

21. Han, M.-J., Kim, B.-Y., Yoon, S.-O., and Chung, A.-S. Cell Proliferation Induced by Reactive Oxygen Species Is Mediated via Mitogen-activated Protein Kinase in Chinese Hamster Lung Fibroblast (V79) Cells

22. Movafagh, S., Crook, S., and Vo, K. (2015) Regulation of hypoxia-inducible Factor-1a by reactive oxygen species: New developments in an old debate. J. Cell. Biochem. 116, 696–703

23. Redza-Dutordoir, M., and Averill-Bates, D. A. (2021) Interactions between reactive oxygen species and autophagy: Special issue: Death mechanisms in cellular homeostasis. Biochim. Biophys. Acta Mol. Cell Res. 10.1016/j.bbamcr.2021.119041

24. Rada, B., Gardina, P., Myers, T. G., and Leto, T. L. (2011) Reactive oxygen species mediate inflammatory cytokine release and EGFR-dependent mucin secretion in airway epithelial cells exposed to Pseudomonas pyocyanin. Mucosal Immunol. 4, 158–171

25. Morgan, M. J., and Liu, Z. G. (2011) Crosstalk of reactive oxygen species and NF-κB signaling. Cell Res. 21, 103–115

26. Lee, Y. W., Lee, W. H., and Kim, P. H. (2010) Oxidative mechanisms of IL-4-induced IL-6 expression in vascular endothelium. Cytokine. 49, 73–79

27. Wruck, C. J., Streetz, K., Pavic, G., Götz, M. E., Tohidnezhad, M., Brandenburg, L. O., Varoga, D., Eickelberg, O., Herdegen, T., Trautwein, C., Cha, K., Kan, Y. W., and Pufe, T. (2011) Nrf2 induces interleukin-6 (IL-6) expression via an antioxidant response element within the IL-6 promoter. Journal of Biological Chemistry. 286, 4493–4499

28. Fan, H., Bai, Q., Yang, Y., Shi, X., Du, G., Yan, J., Shi, J., and Wang, D. (2023) The key roles of reactive oxygen species in microglial inflammatory activation: Regulation by endogenous antioxidant system and exogenous sulfur-containing compounds. Eur. J. Pharmacol. 10.1016/j.ejphar.2023.175966

29. He, F., Ru, X., and Wen, T. (2020) NRF2, a transcription factor for stress response and beyond. Int. J. Mol. Sci. 21, 1–23

30. Montgomery, J. E., Parsons, M. J., and Hyde, D. R. (2010) A novel model of retinal ablation demonstrates that the extent of rod cell death regulates the origin of the regenerated zebrafish rod photoreceptors. Journal of Comparative Neurology. 518, 800–814

31. Boyd, P., and Hyde, D. R. (2022) Iron contributes to photoreceptor degeneration and Müller glia proliferation in the zebrafish light-treated retina. Exp. Eye Res. 10.1016/j.exer.2022.108947

32. Emmerich, K., Walker, S. L., Wang, G., White, D. T., Ceisel, A., Wang, F., Teng, Y., Chunawala, Z., Graziano, G., Nimmagadda, S., Saxena, M. T., Qian, J., and Mumm, J. S. (2023) Transcriptomic comparison of two selective retinal cell ablation paradigms in zebrafish reveals shared and cell-specific regenerative responses. PLoS Genet. 10.1371/journal.pgen.1010905

33. Zhang, H., Yang, S., Lu, Y. L., Zhou, L. Q., Dong, M. H., Chu, Y. H., Pang, X. W., Chen, L., Xu, L. L., Zhang, L. Y., Zhu, L. F., Xu, T., Wang, W., Shang, K., Tian, D. S., and Qin, C. (2025) Microglial Nrf2-mediated lipid and iron metabolism reprogramming promotes remyelination during white matter ischemia. Redox Biol. 10.1016/j.redox.2024.103473

34. Shiau, C. E., Kaufman, Z., Meireles, A. M., and Talbot, W. S. (2015) Differential requirement for irf8 in formation of embryonic and adult macrophages in zebrafish. PLoS One. 10.1371/journal.pone.0117513

35. Mills, M. G., Ramsden, R., Ma, E. Y., Corrales, J., Kristofco, L. A., Steele, W. B., Saari, G. N., Melnikov, F., Kostal, J., Kavanagh, T. J., Zimmerman, J. B., Voutchkova-Kostal, A. M., Brooks, B. W., Coish, P., Anastas, P. T., and Gallagher, E. (2020) CRISPR-Generated Nrf2a Loss- And Gain-of-Function Mutants Facilitate Mechanistic Analysis of Chemical Oxidative Stress-Mediated Toxicity in Zebrafish. Chem. Res. Toxicol. 33, 426–435

36. Lenkowski, J. R., and Raymond, P. A. (2014) Müller glia: Stem cells for generation and regeneration of retinal neurons in teleost fish. Prog. Retin. Eye Res. 40, 94–123

37. Goldman, D. (2014) Müller glial cell reprogramming and retina regeneration. Nat. Rev. Neurosci. 15, 431–442

38. Gao, H., A, L., Huang, X., Chen, X., and Xu, H. (2021) Müller Glia-Mediated Retinal Regeneration. Mol. Neurobiol. 58, 2342–2361

39. Stenkamp, D. L. (2011) The rod photoreceptor lineage of teleost fish. Prog. Retin. Eye Res. 30, 395–404

40. Otteson, D. C., and Hitchcock, P. F. (2003) Stem cells in the teleost retina: Persistent neurogenesis and injury-induced regeneration. in Vision Research, pp. 927–936, Elsevier Ltd, 43, 927–936

41. Lahne, M., Li, J., Marton, R. M., and Hyde, D. R. (2015) Actin-cytoskeleton-and rock-mediated inm are required for photoreceptor regeneration in the adult zebrafish retina. Journal of Neuroscience. 35, 15612–15634

42. Fadool, J. M. (2003) Development of a rod photoreceptor mosaic revealed in transgenic zebrafish. Dev. Biol. 258, 277–290

43. Ching-Hwa Sung, and Andrew W. Tai (1999) Rhodopsin Trafficking and its Role in Retinal Dystrophies. Int. Rev. Cytol. 195, 215–267

44. Reddaway, J., Richardson, P. E., Bevan, R. J., Stoneman, J., and Palombo, M. (2023) Microglial morphometric analysis: so many options, so little consistency. Front. Neuroinform. 10.3389/fninf.2023.1211188

45. Vidal-Itriago, A., Radford, R. A. W., Aramideh, J. A., Maurel, C., Scherer, N. M., Don, E. K., Lee, A., Chung, R. S., Graeber, M. B., and Morsch, M. (2022) Microglia morphophysiological diversity and its implications for the CNS. Front. Immunol. 10.3389/fimmu.2022.997786

46. Blume, Z. I., Lambert, J. M., Lovel, A. G., and Mitchell, D. M. (2020) Microglia in the developing retina couple phagocytosis with the progression of apoptosis via P2RY12 signaling. Developmental Dynamics. 249, 723–740

47. Timme-Laragy, A. R., Karchner, S. I., Franks, D. G., Jenny, M. J., Harbeitner, R. C., Goldstone, J. V., McArthur, A. G., and Hahn, M. E. (2012) Nrf2b, novel zebrafish paralog of oxidant-responsive transcription factor NF-E2-related factor 2 (NRF2). Journal of Biological Chemistry. 287, 4609–4627

48. Katsuragi, Y., Ichimura, Y., and Komatsu, M. (2016) Regulation of the Keap1–Nrf2 pathway by p62/SQSTM1. Curr. Opin. Toxicol. 1, 54–61

49. Jain, A., Lamark, T., Sjøttem, E., Larsen, K. B., Awuh, J. A., Øvervatn, A., McMahon, M., Hayes, J. D., and Johansen, T. (2010) p62/SQSTM1 is a target gene for transcription factor NRF2 and creates a positive feedback loop by inducing antioxidant response element-driven gene transcription. Journal of Biological Chemistry. 285, 22576–22591

50. Bhowmick, S., D’Mello, V., Caruso, D., and Abdul-Muneer, P. M. (2019) Traumatic brain injury-induced downregulation of Nrf2 activates inflammatory response and apoptotic cell death. J. Mol. Med. 97, 1627–1641

51. Zhang, L., Johnson, D., and Johnson, J. A. (2013) Deletion of Nrf2 impairs functional recovery, reduces clearance of myelin debris and decreases axonal remyelination after peripheral nerve injury. Neurobiol. Dis. 54, 329–338

52. Xiong, W., Garfinkel, A. E. M. C., Li, Y., Benowitz, L. I., and Cepko, C. L. (2015) NRF2 promotes neuronal survival in neurodegeneration and acute nerve damage. Journal of Clinical Investigation. 125, 1433–1445

53. Fischer, A. J., Bosse, J. L., and El-Hodiri, H. M. (2013) The ciliary marginal zone (CMZ) in development and regeneration of the vertebrate eye. Exp. Eye Res. 116, 199–204

54. Leach, L. L., Gonzalez, R. G., Jayawardena, S. U., and Gross, J. M. (2025) Macrophage/microglia-dependent mechanisms drive retinal pigment epithelium regeneration in zebrafish. Cell Rep. 10.1016/j.celrep.2025.116325

55. Konar, G. J., Vallone, K. T., Nguyen, T. D., and Patton, J. G. (2025) Analysis of the senescence secretome during zebrafish retina regeneration. Frontiers in Aging. 10.3389/fragi.2025.1569422

56. Reinhard, J., Joachim, S. C., and Faissner, A. (2015) Extracellular matrix remodeling during retinal development. Exp. Eye Res. 133, 132–140

57. Schafer, D. P., Lehrman, E. K., Kautzman, A. G., Koyama, R., Mardinly, A. R., Yamasaki, R., Ransohoff, R. M., Greenberg, M. E., Barres, B. A., and Stevens, B. (2012) Microglia Sculpt Postnatal Neural Circuits in an Activity and Complement-Dependent Manner. Neuron. 74, 691–705

58. Crapser, J. D., Arreola, M. A., Tsourmas, K. I., and Green, K. N. (2021) Microglia as hackers of the matrix: sculpting synapses and the extracellular space. Cell. Mol. Immunol. 18, 2472–2488

59. Harada, T., Harada, C., Kohsaka, S., Wada, E., Yoshida, K., Ohno, S., Mamada, H., Tanaka, K., Parada, L. F., and Wada, K. (2002) Microglia-Mü ller Glia Cell Interactions Control Neurotrophic Factor Production during Light-Induced Retinal Degeneration

60. Ohtsuji, M., Katsuoka, F., Kobayashi, A., Aburatani, H., Hayes, J. D., and Yamamoto, M. (2008) Nrf1 and Nrf2 play distinct roles in activation of antioxidant response element-dependent genes. Journal of Biological Chemistry. 283, 33554–33562

61. Hu, S., Feng, J., Wang, M., Wufuer, R., Liu, K., Zhang, Z., and Zhang, Y. (2022) Nrf1 is an indispensable redox-determining factor for mitochondrial homeostasis by integrating multi-hierarchical regulatory networks. Redox Biol. 10.1016/j.redox.2022.102470

62. Zhou, L. Z. H., Johnson, A. P., and Rando, T. A. (2001) NFkB AND AP-1 MEDIATE TRANSCRIPTIONAL RESPONSES TO OXIDATIVE STRESS IN SKELETAL MUSCLE CELLS. Free Radic. Biol. Med. 31, 1405–1416

63. Lingappan, K. (2018) NF-κB in oxidative stress. Curr. Opin. Toxicol. 7, 81–86

64. Jin, B., Xie, L., Zhan, D., Zhou, L., Feng, Z., He, J., Qin, J., Zhao, C., Luo, L., and Li, L. (2022) Nrf2 dictates the neuronal survival and differentiation of embryonic zebrafish harboring compromised alanyl-tRNA synthetase. Development (Cambridge). 10.1242/dev.200342

65. Sharrock, A. V., Mulligan, T. S., Hall, K. R., Williams, E. M., White, D. T., Zhang, L., Emmerich, K., Matthews, F., Nimmagadda, S., Washington, S., Le, K. D., Meir-Levi, D., Cox, O. L., Saxena, M. T., Calof, A. L., Lopez-Burks, M. E., Lander, A. D., Ding, D., Ji, H., Ackerley, D. F., and Mumm, J. S. (2022) NTR 2.0: a rationally engineered prodrug-converting enzyme with substantially enhanced efficacy for targeted cell ablation. Nat. Methods. 19, 205–215

66. D’Orazi, F. D., Suzuki, S. C., Darling, N., Wong, R. O., and Yoshimatsu, T. (2020) Conditional and biased regeneration of cone photoreceptor types in the zebrafish retina. Journal of Comparative Neurology. 528, 2816–2830

67. Rumford, J. E., Farre, A. A., Mai, J., Weimar, H. V, Shelton, C. D., Morales, M., and Mitchell, D. M. (2026) Cone photoreceptor ablation in microglia-deficient larval zebrafish retina elicits a regenerative response alongside a compensatory immune cell response. BioRxiV. 10.64898/2026.02.26.708140

68. Rumford, J. E., Farre, A. A., Mai, J., Weimar, H. V., Shelton, C. D., Morales, M., and Mitchell, D. M. (2026) Cone photoreceptor ablation in microglia-deficient larval zebrafish retina elicits a regenerative response alongside a compensatory immune cell response. PLoS One. 21, e0344737

69. Thiel, W. A., Blume, Z. I., and Mitchell, D. M. (2022) Compensatory engulfment and Müller glia reactivity in the absence of microglia. Glia. 70, 1402–1425

70. Figueiredo, C., Tellkamp, G., Norden, C., and Rocha-Martins, M. (2026) Apoptosis-induced self-renewal of neurogenic progenitors safeguards retinal development against extensive cell loss. BioRxiV. 10.64898/2026.07.03.735591

71. Conedera, F. M., Pousa, A. M. Q., Mercader, N., Tschopp, M., and Enzmann, V. (2019) Retinal microglia signaling affects Müller cell behavior in the zebrafish following laser injury induction. Glia. 67, 1150–1166

72. Westerfield, M. (2007) The Zebrafish Book. A Guide for the Laboratory Use of Zebrafish (Danio rerio), 5th Ed., University of Oregon Press, Eugene

73. Lovel, A. G., and Mitchell, D. M. (2023) PCNA Staining of Retinal Cryosections to Assess Microglial/Macrophage Proliferation, pp. 389–400, 10.1007/978-1-0716-3012-9_21

74. Meeker, N. D., Hutchinson, S. A., Ho, L., and Trede, N. S. (2007) Method for isolation of PCR-ready genomic DNA from zebrafish tissues. Biotechniques. 43, 610–614

75. McCurley, A. T., and Callard, G. V. (2008) Characterization of housekeeping genes in zebrafish: Male-female differences and effects of tissue type, developmental stage and chemical treatment. BMC Mol. Biol. 10.1186/1471-2199-9-102

76. Howarth, D. L., Lindtner, C., Vacaru, A. M., Sachidanandam, R., Tsedensodnom, O., Vasilkova, T., Buettner, C., and Sadler, K. C. (2014) Activating Transcription Factor 6 Is Necessary and Sufficient for Alcoholic Fatty Liver Disease in Zebrafish. PLoS Genet. 10.1371/journal.pgen.1004335

77. Zhang, R., Varela, M., Vallentgoed, W., Forn-Cuni, G., van der Vaart, M., and Meijer, A. H. (2019) The selective autophagy receptors Optineurin and p62 are both required for zebrafish host resistance to mycobacterial infection. PLoS Pathog. 10.1371/journal.ppat.1007329

