## Supplemental-Information for "Microglia support regeneration of rod photoreceptors in larval zebrafish retina through Nrf2-independent mechanisms"

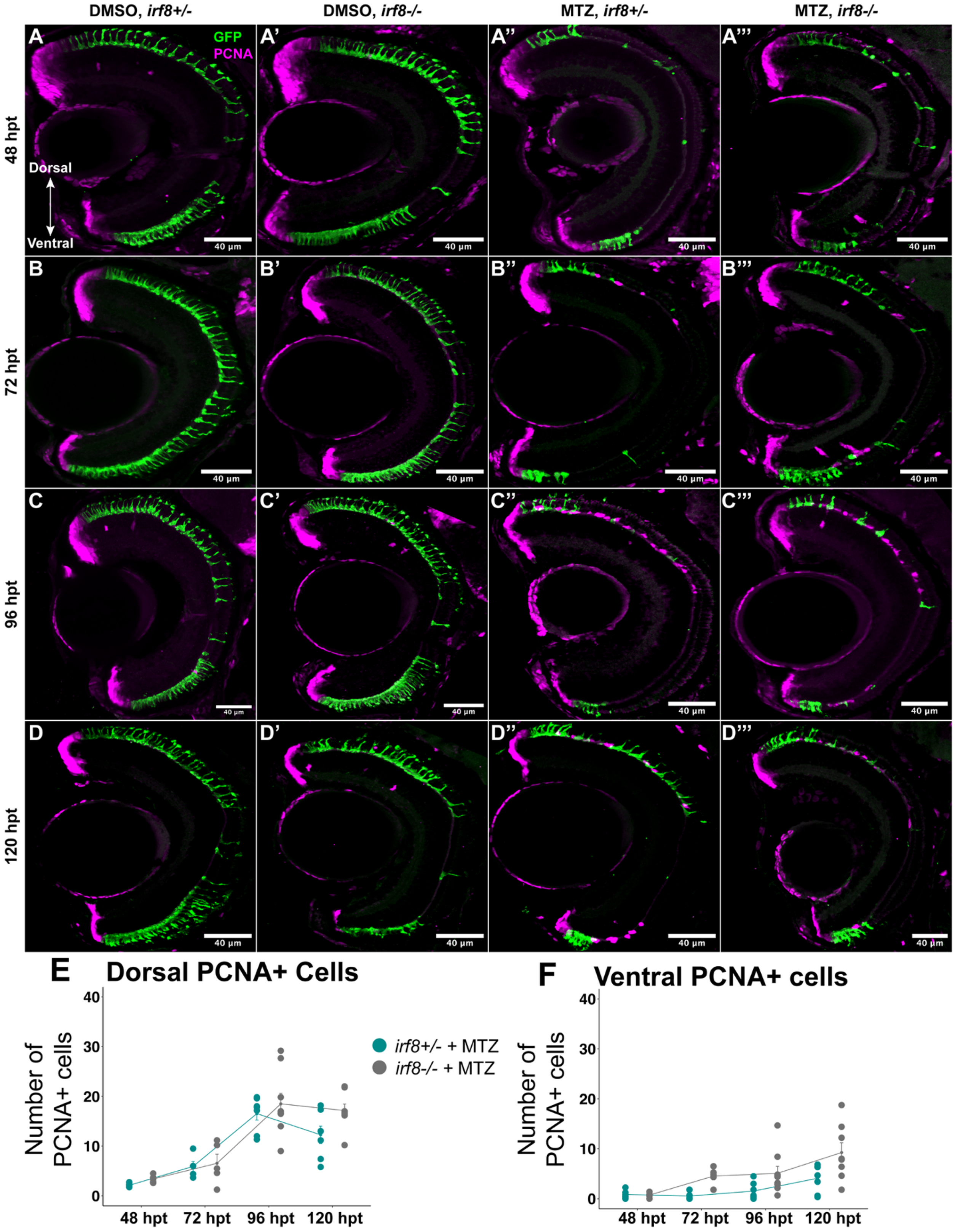

DMSO, irf8+/-
DMSO, irf8-/-
MTZ, irf8+/-
MTZ, irf8-/-
A
A'
A''
A'''
GFP
PCNA
48 hpt
Dorsal
Ventral
40 µm
B
B'
B''
B'''
72 hpt
C
C'
C''
C'''
96 hpt
D
D'
D''
D'''
120 hpt
E
Dorsal PCNA+ Cells
F
Ventral PCNA+ cells
Number of PCNA+ cells
irf8+/- + MTZ
irf8-/- + MTZ
0
10
20
30
40
48 hpt
72 hpt
96 hpt
120 hpt

**Supplemental figure 1: Dorsal and ventral proliferation following rod ablation. A-D''':** Representative images of retinal cryosections from DMSO (control) and MTZ-treated retinas of *irf8+/-* and *irf8-/-* genotypes from 48-120 hpt. Images show PCNA (magenta) and rod cells (green). E-F: Line-graphs showing dorsal (E) and ventral (F) PCNA+ cells.

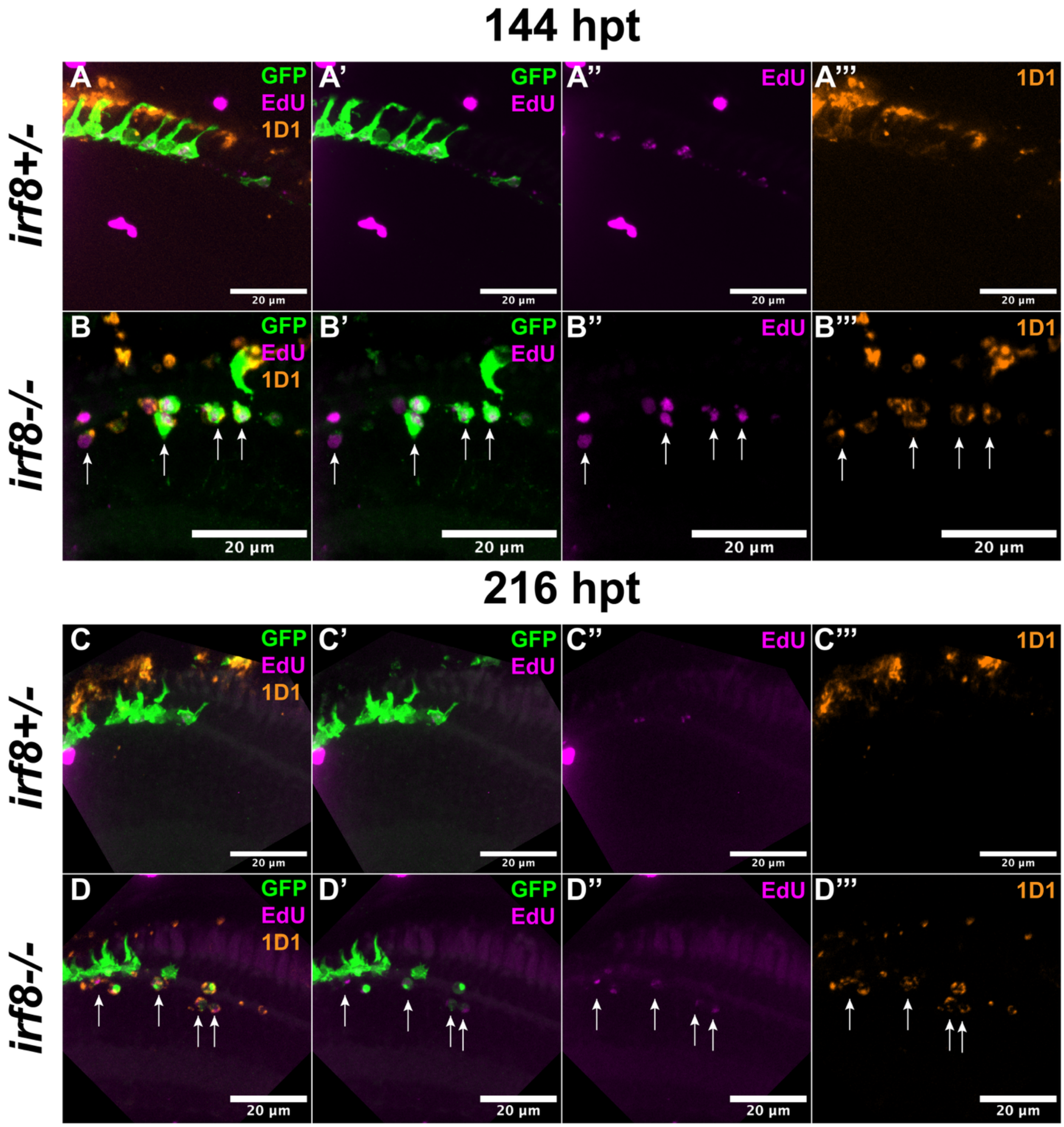

144 hpt
irf8+/-
irf8-/-
A
A'
A''
A'''
B
B'
B''
B'''
GFP
EdU
1D1
20 µm
216 hpt
C
C'
C''
C'''
D
D'
D''
D'''

**Supplemental figure 2: Comparison of ventral regions of MTZ-treated *irf8+/-* and *irf8-/-* retinas at 144- and 216 hpt.** A-B''': Representative images of ventral regions of retinas from MTZ-treated *irf8+/-* (A-A''') and *irf8-/-* (B-B''') larvae at 144 hpt. Images show GFP rods (green), EdU (magenta), and rhodopsin/1D1 (orange). Arrows (seen in B-B''') indicate collapsed EdU+ rod cells. C-D''': Representative images of ventral regions of retinas from MTZ-treated *irf8+/-* (C-C''') and *irf8-/-* (D-D''') larvae at 216 hpt. Images show GFP rods (green), EdU (magenta), and rhodopsin/1D1 (orange). Arrows (seen in D-D''') indicate collapsed EdU+ rod cells.

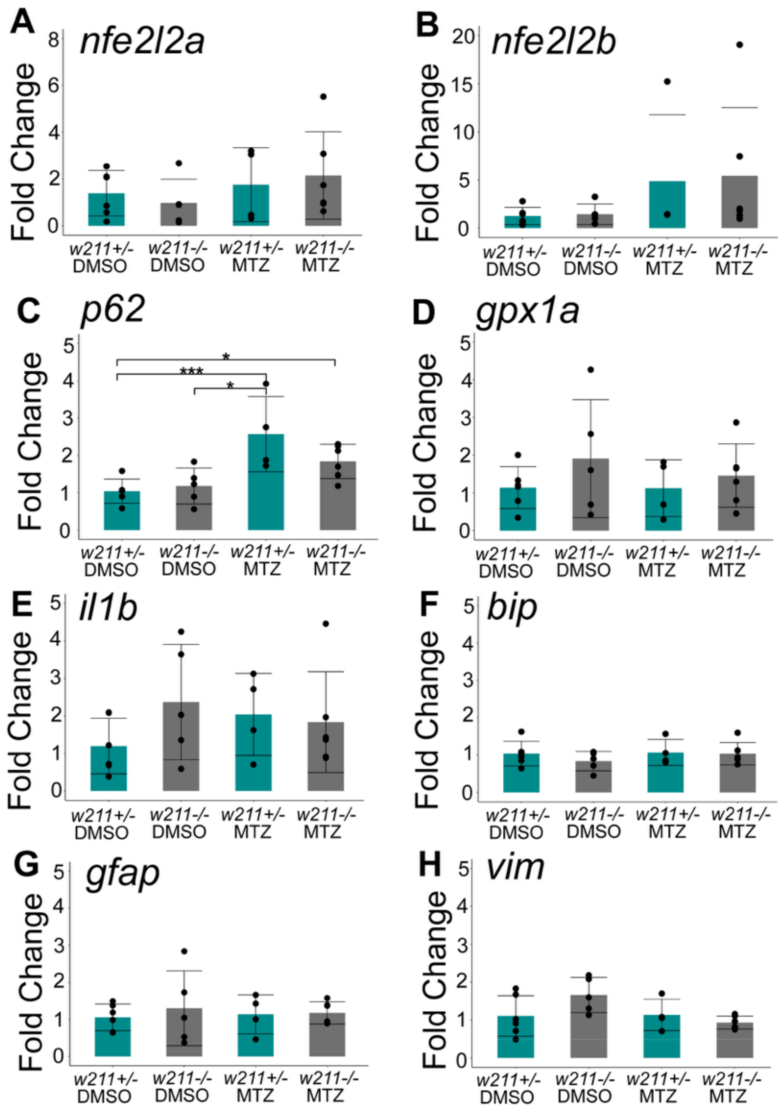


**Supplemental Figure 3: Gene expression measured by RT-qPCR following rod ablation at 48 hpt in *Nrf2*-sufficient and *nrf2* loss-of-function eyes A-H:** Fold change in gene expression measured by RT-qPCR using RNA/cDNA from whole *rho:nfsB-eGFP* larval eyes from the indicated *w211* genotypes, treated with DMSO or MTZ for 48 hpt. * $p<0.05$, ** $p<0.01$, ***$p<0.001$; two-way ANOVA with Tukey's HSD post-hoc test for multiple comparisons. Bars represent mean fold change (ddCt method) relative to DMSO *w211+/-* samples +/- SD, and dots represent biological replicates. For clarity, only statistically significant differences are annotated in these graphs. All p-values are reported in Supporting Information. Sample size: 4-6 larvae per group (both eyes pooled).

### Reporting of actual p-values

### Figure 1:

<u>*48 hpt*</u>

*Total PCNA*

| Comparison (Treatment) | ANOVA p-value |
|---|---|
| MTZ vs DMSO | 0.0083443 |

| Comparison (Genotype) | ANOVA p-value |
|---|---|
| *irf8+/-* vs *irf8-/-* | 0.7754005 |

| Comparison | post-hoc Tukey's HSD |
|---|---|
| DMSO *irf8 +/-* vs MTZ *irf8 +/-* | 0.9216493 |
| DMSO *irf8 +/-* vs DMSO *irf8 -/-* | 0.3535918 |
| DMSO *irf8 +/-* vs MTZ *irf8 -/-* | 0.2582740 |
| DMSO *irf8 -/-* vs MTZ *irf8 +/-* | 0.1320700 |
| MTZ *irf8 -/-* vs MTZ *irf8 +/-* | 0.5796113 |
| DMSO *irf8 -/-* vs MTZ *irf8 -/-* | 0.0109632 |

ONL PCNA

| Comparison (Treatment) | ANOVA p-value |
|---|---|
| MTZ vs DMSO | 0.9334446 |

| Comparison (Genotype) | ANOVA p-value |
|---|---|
| *irf8+/-* vs *irf8-/-* | 0.6165606 |

| Comparison | post-hoc Tukey's HSD |
|---|---|
| DMSO *irf8 +/-* vs MTZ *irf8 +/-* | 0.6795294 |
| DMSO *irf8 +/-* vs DMSO *irf8 -/-* | 0.5016622 |
| DMSO *irf8 +/-* vs MTZ *irf8 -/-* | 0.9740513 |
| DMSO *irf8 -/-* vs MTZ *irf8 +/-* | 0.9901464 |
| MTZ *irf8 -/-* vs MTZ *irf8 +/-* | 0.8937555 |
| DMSO *irf8 -/-* vs MTZ *irf8 -/-* | 0.7483535 |

INL PCNA

| Comparison (Treatment) | ANOVA p-value |
|---|---|
| MTZ vs DMSO | 0.0016208 |

| Comparison (Genotype) | ANOVA p-value |
|---|---|
| *irf8+/-* vs *irf8-/-* | 0.9613959 |

| Comparison | post-hoc Tukey's HSD |
|---|---|
| DMSO *irf8 +/-* vs MTZ *irf8 +/-* | 0.5634961 |
| DMSO *irf8 +/-* vs DMSO *irf8 -/-* | 0.5226395 |
| DMSO *irf8 +/-* vs MTZ *irf8 -/-* | 0.0755753 |
| DMSO *irf8 -/-* vs MTZ *irf8 +/-* | 0.0663789 |
| MTZ *irf8 -/-* vs MTZ *irf8 +/-* | 0.5634961 |
| DMSO *irf8 -/-* vs MTZ *irf8 -/-* | 0.0047923 |

PCNA+ Müller glia cells

| Comparison (Treatment) | ANOVA p-value |
|---|---|
| MTZ vs DMSO | 0.040594 |

| Comparison (Genotype) | ANOVA p-value |
|---|---|

| | |
|---|---|
| *irf8+/-* vs *irf8-/-* | 0.6763894 |

| Comparison | post-hoc Tukey's HSD |
|---|---|
| DMSO *irf8 +/-* vs MTZ *irf8 +/-* | 0.9780938 |
| DMSO *irf8 +/-* vs DMSO *irf8 -/-* | 0.1462666 |
| DMSO *irf8 +/-* vs MTZ *irf8 -/-* | 0.5910768 |
| DMSO *irf8 -/-* vs MTZ *irf8 +/-* | 0.2766289 |
| MTZ *irf8 -/-* vs MTZ *irf8 +/-* | 0.3697008 |
| DMSO *irf8 -/-* vs MTZ *irf8 -/-* | 0.0128511 |

Rod cells

| Comparison (Treatment) | ANOVA p-value |
|---|---|
| MTZ vs DMSO | 5e-06 |

| Comparison (Genotype) | ANOVA p-value |
|---|---|
| *irf8+/-* vs *irf8-/-* | 0.219488 |

| Comparison | post-hoc Tukey's HSD |
|---|---|
| DMSO *irf8 +/-* vs MTZ *irf8 +/-* | 0.0344425 |
| DMSO *irf8 +/-* vs DMSO *irf8 -/-* | 0.0819284 |
| DMSO *irf8 +/-* vs MTZ *irf8 -/-* | 0.0070326 |
| DMSO *irf8 -/-* vs MTZ *irf8 +/-* | 0.0001914 |
| MTZ *irf8 -/-* vs MTZ *irf8 +/-* | 0.8574034 |
| DMSO *irf8 -/-* vs MTZ *irf8 -/-* | 0.0000438 |

*72 hpt*

Total PCNA

| Comparison (Treatment) | ANOVA p-value |
|---|---|

| | |
|---|---|
| MTZ vs DMSO | 9.9e-06 |

| Comparison (Genotype) | ANOVA p-value |
|---|---|
| *irf8*+/- vs *irf8*-/- | 0.3448609 |

| Comparison | post-hoc Tukey's HSD |
|---|---|
| DMSO *irf8* +/- vs MTZ *irf8* +/- | 0.1390951 |
| DMSO *irf8* +/- vs DMSO *irf8* -/- | 0.6372465 |
| DMSO *irf8* +/- vs MTZ *irf8* -/- | 0.0003550 |
| DMSO *irf8* -/- vs MTZ *irf8* +/- | 0.0146116 |
| MTZ *irf8* -/- vs MTZ *irf8* +/- | 0.0582805 |
| DMSO *irf8* -/- vs MTZ *irf8* -/- | 0.0000340 |

ONL PCNA

| Comparison (Treatment) | ANOVA p-value |
|---|---|
| MTZ vs DMSO | 7.7e-06 |

| Comparison (Genotype) | ANOVA p-value |
|---|---|
| *irf8*+/- vs *irf8*-/- | 0.6384757 |

| Comparison | post-hoc Tukey's HSD |
|---|---|
| DMSO *irf8* +/- vs MTZ *irf8* +/- | 0.0929666 |
| DMSO *irf8* +/- vs DMSO *irf8* -/- | 0.1797904 |
| DMSO *irf8* +/- vs MTZ *irf8* -/- | 0.0027114 |
| DMSO *irf8* -/- vs MTZ *irf8* +/- | 0.0013196 |
| MTZ *irf8* -/- vs MTZ *irf8* +/- | 0.3875702 |
| DMSO *irf8* -/- vs MTZ *irf8* -/- | 0.0000376 |

INL PCNA

| Comparison (Treatment) | ANOVA p-value |
|---|---|
| MTZ vs DMSO | 0.0044952 |

| Comparison (Genotype) | ANOVA p-value |
|---|---|
| *irf8*+/- vs *irf8*-/- | 0.0153122 |

| Comparison | post-hoc Tukey's HSD |
|---|---|
| DMSO *irf8* +/- vs MTZ *irf8* +/- | 0.8350767 |
| DMSO *irf8* +/- vs DMSO *irf8* -/- | 0.9334096 |
| DMSO *irf8* +/- vs MTZ *irf8* -/- | 0.0021574 |
| DMSO *irf8* -/- vs MTZ *irf8* +/- | 0.9924284 |
| MTZ *irf8* -/- vs MTZ *irf8* +/- | 0.0179507 |
| DMSO *irf8* -/- vs MTZ *irf8* -/- | 0.0072902 |

PCNA+ Müller glia cells

| Comparison (Treatment) | ANOVA p-value |
|---|---|
| MTZ vs DMSO | 0.0506339 |

| Comparison (Genotype) | ANOVA p-value |
|---|---|
| *irf8*+/- vs *irf8*-/- | 0.0898988 |

| Comparison | post-hoc Tukey's HSD |
|---|---|
| DMSO *irf8* +/- vs MTZ *irf8* +/- | 0.9797300 |
| DMSO *irf8* +/- vs DMSO *irf8* -/- | 0.9920362 |
| DMSO *irf8* +/- vs MTZ *irf8* -/- | 0.0483991 |
| DMSO *irf8* -/- vs MTZ *irf8* +/- | 0.9993675 |
| MTZ *irf8* -/- vs MTZ *irf8* +/- | 0.1232935 |
| DMSO *irf8* -/- vs MTZ *irf8* -/- | 0.0814467 |

Rod cells

| Comparison (Treatment) | ANOVA p-value |
|---|---|
| MTZ vs DMSO | 4.9e-06 |

| Comparison (Genotype) | ANOVA p-value |
|---|---|
| *irf8+/-* vs *irf8-/-* | 0.8790417 |

| Comparison | post-hoc Tukey's HSD |
|---|---|
| DMSO *irf8 +/-* vs MTZ *irf8 +/-* | 0.0254220 |
| DMSO *irf8 +/-* vs DMSO *irf8 -/-* | 0.6370480 |
| DMSO *irf8 +/-* vs MTZ *irf8 -/-* | 0.0008116 |
| DMSO *irf8 -/-* vs MTZ *irf8 +/-* | 0.0022226 |
| MTZ *irf8 -/-* vs MTZ *irf8 +/-* | 0.4361919 |
| DMSO *irf8 -/-* vs MTZ *irf8 -/-* | 0.0000742 |

*96 hpt*

Total PCNA

| Comparison (Treatment) | ANOVA p-value |
|---|---|
| MTZ vs DMSO | 2e-07 |

| Comparison (Genotype) | ANOVA p-value |
|---|---|
| *irf8+/-* vs *irf8-/-* | 0.6859006 |

| Comparison | post-hoc Tukey's HSD |
|---|---|
| DMSO *irf8 +/-* vs MTZ *irf8 +/-* | 0.0023206 |
| DMSO *irf8 +/-* vs DMSO *irf8 -/-* | 0.9442468 |

| | |
|---|---|
| DMSO *irf8* +/- vs MTZ *irf8* -/- | 0.0000989 |
| DMSO *irf8* -/- vs MTZ *irf8* +/- | 0.0005685 |
| MTZ *irf8* -/- vs MTZ *irf8* +/- | 0.6883672 |
| DMSO *irf8* -/- vs MTZ *irf8* -/- | 0.0000232 |

ONL PCNA

| Comparison (Treatment) | ANOVA p-value |
|---|---|
| MTZ vs DMSO | 2e-07 |

| Comparison (Genotype) | ANOVA p-value |
|---|---|
| *irf8*+/- vs *irf8*-/- | 0.5089654 |

| Comparison | post-hoc Tukey's HSD |
|---|---|
| DMSO *irf8* +/- vs MTZ *irf8* +/- | 0.0022428 |
| DMSO *irf8* +/- vs DMSO *irf8* -/- | 0.9778549 |
| DMSO *irf8* +/- vs MTZ *irf8* -/- | 0.0000555 |
| DMSO *irf8* -/- vs MTZ *irf8* +/- | 0.0008155 |
| MTZ *irf8* -/- vs MTZ *irf8* +/- | 0.5589807 |
| DMSO *irf8* -/- vs MTZ *irf8* -/- | 0.0000196 |

INL PCNA

| Comparison (Treatment) | ANOVA p-value |
|---|---|
| MTZ vs DMSO | 0.5628259 |

| Comparison (Genotype) | ANOVA p-value |
|---|---|
| *irf8*+/- vs *irf8*-/- | 0.0100769 |

| Comparison | post-hoc Tukey's HSD |
|---|---|

| | |
|---|---|
| DMSO *irf8 +/-* vs MTZ *irf8 +/-* | 0.8041108 |
| DMSO *irf8 +/-* vs DMSO *irf8 -/-* | 0.8987497 |
| DMSO *irf8 +/-* vs MTZ *irf8 -/-* | 0.1310488 |
| DMSO *irf8 -/-* vs MTZ *irf8 +/-* | 0.3996785 |
| MTZ *irf8 -/-* vs MTZ *irf8 +/-* | 0.0179683 |
| DMSO *irf8 -/-* vs MTZ *irf8 -/-* | 0.4189268 |

PCNA+ Müller glia cells

| Comparison (Treatment) | ANOVA p-value |
|---|---|
| MTZ vs DMSO | 0.7828939 |

| Comparison (Genotype) | ANOVA p-value |
|---|---|
| *irf8+/-* vs *irf8-/-* | 0.004711 |

| Comparison | post-hoc Tukey's HSD |
|---|---|
| DMSO *irf8 +/-* vs MTZ *irf8 +/-* | 0.7728295 |
| DMSO *irf8 +/-* vs DMSO *irf8 -/-* | 0.7060508 |
| DMSO *irf8 +/-* vs MTZ *irf8 -/-* | 0.1284034 |
| DMSO *irf8 -/-* vs MTZ *irf8 +/-* | 0.2013816 |
| MTZ *irf8 -/-* vs MTZ *irf8 +/-* | 0.0152029 |
| DMSO *irf8 -/-* vs MTZ *irf8 -/-* | 0.6488137 |

Rod cells

| Comparison (Treatment) | ANOVA p-value |
|---|---|
| MTZ vs DMSO | 2e-07 |

| Comparison (Genotype) | ANOVA p-value |
|---|---|
| *irf8+/-* vs *irf8-/-* | 0.7339297 |

| Comparison | post-hoc Tukey's HSD |
| --- | --- |
| DMSO *irf8* +/- vs MTZ *irf8* +/- | 0.0027255 |
| DMSO *irf8* +/- vs DMSO *irf8* -/- | 0.6345865 |
| DMSO *irf8* +/- vs MTZ *irf8* -/- | 0.0003490 |
| DMSO *irf8* -/- vs MTZ *irf8* +/- | 0.0001302 |
| MTZ *irf8* -/- vs MTZ *irf8* +/- | 0.9037414 |
| DMSO *irf8* -/- vs MTZ *irf8* -/- | 0.0000152 |

*120 hpt*

Total PCNA

| Comparison (Treatment) | ANOVA p-value |
| --- | --- |
| MTZ vs DMSO | 1e-07 |

| Comparison (Genotype) | ANOVA p-value |
| --- | --- |
| *irf8+/-* vs *irf8-/-* | 0.0260935 |

| Comparison | post-hoc Tukey's HSD |
| --- | --- |
| DMSO *irf8* +/- vs MTZ *irf8* +/- | 0.0048411 |
| DMSO *irf8* +/- vs DMSO *irf8* -/- | 0.9967787 |
| DMSO *irf8* +/- vs MTZ *irf8* -/- | 0.0000054 |
| DMSO *irf8* -/- vs MTZ *irf8* +/- | 0.0034972 |
| MTZ *irf8* -/- vs MTZ *irf8* +/- | 0.0305825 |
| DMSO *irf8* -/- vs MTZ *irf8* -/- | 0.0000020 |

ONL PCNA

| Comparison (Treatment) | ANOVA p-value |
| --- | --- |
| MTZ vs DMSO | 4e-07 |

| Comparison (Genotype) | ANOVA p-value |
|---|---|
| *irf8+/-* vs *irf8-/-* | 0.1967037 |

| Comparison | post-hoc Tukey's HSD |
|---|---|
| DMSO *irf8 +/-* vs MTZ *irf8 +/-* | 0.0048949 |
| DMSO *irf8 +/-* vs DMSO *irf8 -/-* | 0.9999828 |
| DMSO *irf8 +/-* vs MTZ *irf8 -/-* | 0.0000700 |
| DMSO *irf8 -/-* vs MTZ *irf8 +/-* | 0.0018591 |
| MTZ *irf8 -/-* vs MTZ *irf8 +/-* | 0.2950374 |
| DMSO *irf8 -/-* vs MTZ *irf8 -/-* | 0.0000166 |

INL PCNA

| Comparison (Treatment) | ANOVA p-value |
|---|---|
| MTZ vs DMSO | 0.0287214 |

| Comparison (Genotype) | ANOVA p-value |
|---|---|
| *irf8+/-* vs *irf8-/-* | 0.0416161 |

| Comparison | post-hoc Tukey's HSD |
|---|---|
| DMSO *irf8 +/-* vs MTZ *irf8 +/-* | 0.9771535 |
| DMSO *irf8 +/-* vs DMSO *irf8 -/-* | 0.9961345 |
| DMSO *irf8 +/-* vs MTZ *irf8 -/-* | 0.0413486 |
| DMSO *irf8 -/-* vs MTZ *irf8 +/-* | 0.9970084 |
| MTZ *irf8 -/-* vs MTZ *irf8 +/-* | 0.0588918 |
| DMSO *irf8 -/-* vs MTZ *irf8 -/-* | 0.0378388 |

PCNA+ Müller glia cells

| Comparison (Treatment) | ANOVA p-value |
|---|---|
| MTZ vs DMSO | 0.0120694 |

| Comparison (Genotype) | ANOVA p-value |
|---|---|
| *irf8*+/- vs *irf8*-/- | 0.0380441 |

| Comparison | post-hoc Tukey's HSD |
|---|---|
| DMSO *irf8* +/- vs MTZ *irf8* +/- | 0.6769378 |
| DMSO *irf8* +/- vs DMSO *irf8* -/- | 0.8916652 |
| DMSO *irf8* +/- vs MTZ *irf8* -/- | 0.0169877 |
| DMSO *irf8* -/- vs MTZ *irf8* +/- | 0.9678268 |
| MTZ *irf8* -/- vs MTZ *irf8* +/- | 0.1243570 |
| DMSO *irf8* -/- vs MTZ *irf8* -/- | 0.0483343 |

Rod cells

| Comparison (Treatment) | ANOVA p-value |
|---|---|
| MTZ vs DMSO | 5.7e-06 |

| Comparison (Genotype) | ANOVA p-value |
|---|---|
| *irf8*+/- vs *irf8*-/- | 0.9778856 |

| Comparison | post-hoc Tukey's HSD |
|---|---|
| DMSO *irf8* +/- vs MTZ *irf8* +/- | 0.0224882 |
| DMSO *irf8* +/- vs DMSO *irf8* -/- | 0.8709836 |
| DMSO *irf8* +/- vs MTZ *irf8* -/- | 0.0048197 |
| DMSO *irf8* -/- vs MTZ *irf8* +/- | 0.0015425 |
| MTZ *irf8* -/- vs MTZ *irf8* +/- | 0.9195447 |
| DMSO *irf8* -/- vs MTZ *irf8* -/- | 0.0002314 |

### Figure 2:

*Total EdU+ rods*

| Comparison (Treatment) | ANOVA p-value |
|---|---|
| MTZ vs DMSO | 0 |

| Comparison (Genotype) | ANOVA p-value |
|---|---|
| *irf8*+/- vs *irf8*-/- | 0.0081769 |

| Comparison | post-hoc Tukey's HSD |
|---|---|
| DMSO *irf8* +/- vs MTZ *irf8* +/- | 0.0000080 |
| DMSO *irf8* +/- vs DMSO *irf8* -/- | 0.8262375 |
| DMSO *irf8* +/- vs MTZ *irf8* -/- | 0.0662517 |
| DMSO *irf8* -/- vs MTZ *irf8* +/- | 0.0000001 |
| MTZ *irf8* -/- vs MTZ *irf8* +/- | 0.0155291 |
| DMSO *irf8* -/- vs MTZ *irf8* -/- | 0.0058625 |

*Dorsal EdU+ rods*

| Comparison (Treatment) | ANOVA p-value |
|---|---|
| MTZ vs DMSO | 0 |

| Comparison (Genotype) | ANOVA p-value |
|---|---|
| *irf8*+/- vs *irf8*-/- | 0.0032508 |

| Comparison | post-hoc Tukey's HSD |
|---|---|
| DMSO *irf8* +/- vs MTZ *irf8* +/- | 0.0000049 |
| DMSO *irf8* +/- vs DMSO *irf8* -/- | 0.7063635 |
| DMSO *irf8* +/- vs MTZ *irf8* -/- | 0.0843450 |

| | |
|---|---|
| DMSO *irf8 -/-* vs MTZ *irf8 +/-* | 0.0000000 |
| MTZ *irf8 -/-* vs MTZ *irf8 +/-* | 0.0069135 |
| DMSO *irf8 -/-* vs MTZ *irf8 -/-* | 0.0046951 |

*Ventral EdU+ rods*

| Comparison | Conover test with Bonferroni correction |
|---|---|
| DMSO *irf8 +/-* vs MTZ *irf8 +/-* | 0.0001 |
| DMSO *irf8 +/-* vs DMSO *irf8 -/-* | 1.0000 |
| DMSO *irf8 +/-* vs MTZ *irf8 -/-* | 0.0059 |
| DMSO *irf8 -/-* vs MTZ *irf8 +/-* | 0.0000 |
| MTZ *irf8 -/-* vs MTZ *irf8 +/-* | 0.9998 |
| DMSO *irf8 -/-* vs MTZ *irf8 -/-* | 0.0030 |

*Total rods*

| Comparison (Treatment) | ANOVA p-value |
|---|---|
| MTZ vs DMSO | 8e-07 |

| Comparison (Genotype) | ANOVA p-value |
|---|---|
| *irf8+/-* vs *irf8-/-* | 0.5913972 |

| Comparison | post-hoc Tukey's HSD |
|---|---|
| DMSO *irf8 +/-* vs MTZ *irf8 +/-* | 0.0000370 |
| DMSO *irf8 +/-* vs DMSO *irf8 -/-* | 0.5026738 |
| DMSO *irf8 +/-* vs MTZ *irf8 -/-* | 0.0009443 |
| DMSO *irf8 -/-* vs MTZ *irf8 +/-* | 0.0005065 |
| MTZ *irf8 -/-* vs MTZ *irf8 +/-* | 0.9496944 |
| DMSO *irf8 -/-* vs MTZ *irf8 -/-* | 0.0125171 |

*Total dorsal rods*

| Comparison (Treatment) | ANOVA p-value |
|---|---|
| MTZ vs DMSO | 2.6e-06 |

| Comparison (Genotype) | ANOVA p-value |
|---|---|
| *irf8*+/- vs *irf8*-/- | 0.387758 |

| Comparison | post-hoc Tukey's HSD |
|---|---|
| DMSO *irf8* +/- vs MTZ *irf8* +/- | 0.0003679 |
| DMSO *irf8* +/- vs DMSO *irf8* -/- | 0.7786368 |
| DMSO *irf8* +/- vs MTZ *irf8* -/- | 0.0009869 |
| DMSO *irf8* -/- vs MTZ *irf8* +/- | 0.0016219 |
| MTZ *irf8* -/- vs MTZ *irf8* +/- | 0.9826049 |
| DMSO *irf8* -/- vs MTZ *irf8* -/- | 0.0043004 |

*Total ventral rods*

| Comparison (Treatment) | ANOVA p-value |
|---|---|
| MTZ vs DMSO | 0.0003308 |

| Comparison (Genotype) | ANOVA p-value |
|---|---|
| *irf8*+/- vs *irf8*-/- | 0.1999027 |

| Comparison | post-hoc Tukey's HSD |
|---|---|
| DMSO *irf8* +/- vs MTZ *irf8* +/- | 0.0001269 |
| DMSO *irf8* +/- vs DMSO *irf8* -/- | 0.0235299 |
| DMSO *irf8* +/- vs MTZ *irf8* -/- | 0.0070233 |
| DMSO *irf8* -/- vs MTZ *irf8* +/- | 0.1860794 |
| MTZ *irf8* -/- vs MTZ *irf8* +/- | 0.7538510 |

| | |
|---|---|
| DMSO *irf8 -/-* vs MTZ *irf8 -/-* | 0.8506127 |

*Collapsed EdU+ rods*

| Comparison (Genotype) | U-test p-value |
|---|---|
| *irf8+/-* vs *irf8-/-* | 0.003783 |

*Total Collapsed + Rods*

| Comparison (Genotype) | U-test p-value |
|---|---|
| *irf8+/-* vs *irf8-/-* | 0.00075 |

### Figure 3:

*Total EdU+ rods*

| Comparison (Treatment) | ANOVA p-value |
|---|---|
| MTZ vs DMSO | 0 |

| Comparison (Genotype) | ANOVA p-value |
|---|---|
| *irf8+/-* vs *irf8-/-* | 0.4402592 |

| Comparison | post-hoc Tukey's HSD |
|---|---|
| DMSO *irf8 +/-* vs MTZ *irf8 +/-* | 0.0000601 |
| DMSO *irf8 +/-* vs DMSO *irf8 -/-* | 0.9994955 |
| DMSO *irf8 +/-* vs MTZ *irf8 -/-* | 0.0002467 |
| DMSO *irf8 -/-* vs MTZ *irf8 +/-* | 0.0000258 |
| MTZ *irf8 -/-* vs MTZ *irf8 +/-* | 0.6073688 |

| | |
|---|---|
| DMSO *irf8 -/-* vs MTZ *irf8 -/-* | 0.0000847 |

*Dorsal EdU+ rods*

| Comparison (Treatment) | ANOVA p-value |
|---|---|
| MTZ vs DMSO | 2e-07 |

| Comparison (Genotype) | ANOVA p-value |
|---|---|
| *irf8+/-* vs *irf8-/-* | 0.4245478 |

| Comparison | post-hoc Tukey's HSD |
|---|---|
| DMSO *irf8 +/-* vs MTZ *irf8 +/-* | 0.0002204 |
| DMSO *irf8 +/-* vs DMSO *irf8 -/-* | 0.9999802 |
| DMSO *irf8 +/-* vs MTZ *irf8 -/-* | 0.0009719 |
| DMSO *irf8 -/-* vs MTZ *irf8 +/-* | 0.0000862 |
| MTZ *irf8 -/-* vs MTZ *irf8 +/-* | 0.6296547 |
| DMSO *irf8 -/-* vs MTZ *irf8 -/-* | 0.0003114 |

*Ventral EdU+ rods*

| Comparison | Conover test with Bonferroni correction |
|---|---|
| DMSO *irf8 +/-* vs MTZ *irf8 +/-* | 0.0000 |
| DMSO *irf8 +/-* vs DMSO *irf8 -/-* | 1.0000 |
| DMSO *irf8 +/-* vs MTZ *irf8 -/-* | 0.0000 |
| DMSO *irf8 -/-* vs MTZ *irf8 +/-* | 0.0000 |
| MTZ *irf8 -/-* vs MTZ *irf8 +/-* | 0.5666 |
| DMSO *irf8 -/-* vs MTZ *irf8 -/-* | 0.0000 |

*Total rods*

| Comparison (Treatment) | ANOVA p-value |
| --- | --- |
| MTZ vs DMSO | 0 |

| Comparison (Genotype) | ANOVA p-value |
| --- | --- |
| *irf8*+/- vs *irf8*-/- | 0.5809616 |

| Comparison | post-hoc Tukey's HSD |
| --- | --- |
| DMSO *irf8* +/- vs MTZ *irf8* +/- | 0.0191209 |
| DMSO *irf8* +/- vs DMSO *irf8* -/- | 0.5252549 |
| DMSO *irf8* +/- vs MTZ *irf8* -/- | 0.0000097 |
| DMSO *irf8* -/- vs MTZ *irf8* +/- | 0.0003158 |
| MTZ *irf8* -/- vs MTZ *irf8* +/- | 0.1417404 |
| DMSO *irf8* -/- vs MTZ *irf8* -/- | 0.0000000 |

*Total dorsal rods*

| Comparison (Treatment) | ANOVA p-value |
| --- | --- |
| MTZ vs DMSO | 0 |

| Comparison (Genotype) | ANOVA p-value |
| --- | --- |
| *irf8*+/- vs *irf8*-/- | 0.6913955 |

| Comparison | post-hoc Tukey's HSD |
| --- | --- |
| DMSO *irf8* +/- vs MTZ *irf8* +/- | 0.0113437 |
| DMSO *irf8* +/- vs DMSO *irf8* -/- | 0.7062142 |
| DMSO *irf8* +/- vs MTZ *irf8* -/- | 0.0000234 |
| DMSO *irf8* -/- vs MTZ *irf8* +/- | 0.0003679 |
| MTZ *irf8* -/- vs MTZ *irf8* +/- | 0.3489603 |
| DMSO *irf8* -/- vs MTZ *irf8* -/- | 0.0000002 |

*Total ventral rods*

| Comparison (Treatment) | ANOVA p-value |
| --- | --- |
| MTZ vs DMSO | 0 |

| Comparison (Genotype) | ANOVA p-value |
| --- | --- |
| *irf8+/-* vs *irf8-/-* | 0.8570177 |

| Comparison | post-hoc Tukey's HSD |
| --- | --- |
| DMSO *irf8 +/-* vs MTZ *irf8 +/-* | 0.0101391 |
| DMSO *irf8 +/-* vs DMSO *irf8 -/-* | 0.5977667 |
| DMSO *irf8 +/-* vs MTZ *irf8 -/-* | 0.0000298 |
| DMSO *irf8 -/-* vs MTZ *irf8 +/-* | 0.0002005 |
| MTZ *irf8 -/-* vs MTZ *irf8 +/-* | 0.4194042 |
| DMSO *irf8 -/-* vs MTZ *irf8 -/-* | 0.0000001 |

*Collapsed EdU+ rods*

| Comparison (Genotype) | U-test p-value |
| --- | --- |
| *irf8+/-* vs *irf8-/-* | 0.002309 |

*Total Collapsed + Rods*

| Comparison (Genotype) | U-test p-value |
| --- | --- |
| *irf8+/-* vs *irf8-/-* | 0.002415 |

### Figure 4:

*Collapsed EdU+ rods (all MTZ)*

| Comparison | Conover test with Bonferroni correction |
|---|---|
| 144 hpt *irf8*+/- vs 216 hpt *irf8* +/- | 0.3278 |
| 144 hpt *irf8* +/- vs 144 hpt *irf8* -/- | 0.0000 |
| 216 hpt *irf8* +/- vs 144 hpt *irf8* -/- | 0.0000 |
| 216 hpt *irf8* -/- vs 144 hpt *irf8* +/- | 0.0335 |
| 216 hpt *irf8* -/- vs 216 hpt *irf8* +/- | 0.0003 |
| 216 hpt *irf8* -/- vs 144 hpt *irf8* -/- | 0.0226 |

*Collapsed EdU+ rods (all MTZ)*

| Comparison | Conover test with Bonferroni correction |
|---|---|
| 144 hpt *irf8*+/- vs 216 hpt *irf8* +/- | 0.0031 |
| 144 hpt *irf8* +/- vs 144 hpt *irf8* -/- | 0.0000 |
| 216 hpt *irf8* +/- vs 144 hpt *irf8* -/- | 0.0000 |
| 216 hpt *irf8* -/- vs 144 hpt *irf8* +/- | 0.0009 |
| 216 hpt *irf8* -/- vs 216 hpt *irf8* +/- | 0.0000 |
| 216 hpt *irf8* -/- vs 144 hpt *irf8* -/- | 0.2381 |

### Figure 5:

*Corrected total cell fluorescence*

| Comparison (Treatment) | ANOVA p-value |
|---|---|
| MTZ vs DMSO | 0 |

| Comparison (Genotype) | ANOVA p-value |
|---|---|
| *w211+/-* vs *w211-/-* | 0.3891175 |

| Comparison | post-hoc Tukey's HSD |
|---|---|
| DMSO *w211 +/-* vs MTZ *w211 +/-* | 0.0000000 |
| DMSO *w211 +/-* vs DMSO *w211 -/-* | 0.0108476 |
| DMSO *w211 +/-* vs MTZ *w211 -/-* | 0.0000000 |
| DMSO *w211 -/-* vs MTZ *w211 +/-* | 0.0000000 |
| MTZ *w211 -/-* vs MTZ *w211 +/-* | 0.0019151 |
| DMSO *w211 -/-* vs MTZ *w211 -/-* | 0.0000000 |

*Microglia*

| Comparison (Treatment) | ANOVA p-value |
|---|---|
| MTZ vs DMSO | 0.0001968 |

| Comparison (Genotype) | ANOVA p-value |
|---|---|
| *w211+/-* vs *w211-/-* | 0.934138 |

| Comparison | post-hoc Tukey's HSD |
|---|---|
| DMSO *w211 +/-* vs MTZ *w211 +/-* | 0.0200363 |
| DMSO *w211 +/-* vs DMSO *w211 -/-* | 0.9972163 |
| DMSO *w211 +/-* vs MTZ *w211 -/-* | 0.0148588 |
| DMSO *w211 -/-* vs MTZ *w211 +/-* | 0.0383069 |
| MTZ *w211 -/-* vs MTZ *w211 +/-* | 0.9999649 |
| DMSO *w211 -/-* vs MTZ *w211 -/-* | 0.0312941 |

*Activated morphology*

| Comparison (Treatment) | ANOVA p-value |
|---|---|
| MTZ vs DMSO | 3e-07 |

| Comparison (Genotype) | ANOVA p-value |
|---|---|
| *w211*+/- vs *w211*-/- | 0.1114347 |

| Comparison | post-hoc Tukey's HSD |
|---|---|
| DMSO *w211* +/- vs MTZ *w211* +/- | 0.0008037 |
| DMSO *w211* +/- vs DMSO *w211* -/- | 0.9810831 |
| DMSO *w211* +/- vs MTZ *w211* -/- | 0.0000077 |
| DMSO *w211* -/- vs MTZ *w211* +/- | 0.0113911 |
| MTZ *w211* -/- vs MTZ *w211* +/- | 0.2862979 |
| DMSO *w211* -/- vs MTZ *w211* -/- | 0.0002560 |

*Collapsed morphology*

(Non-parametric) Conover test with Bonferroni correction

| Comparison | p-value |
|---|---|
| DMSO *w211* +/- vs MTZ *w211* +/- | 0.0128 |
| DMSO *w211* +/- vs DMSO *w211* -/- | 1.0000 |
| DMSO *w211* +/- vs MTZ *w211* -/- | 0.0182 |
| DMSO *w211* -/- vs MTZ *w211* +/- | 0.2507 |
| MTZ *w211* -/- vs MTZ *w211* +/- | 1.0000 |
| DMSO *w211* -/- vs MTZ *w211* -/- | 0.3386 |

*ONL/OPL-localized microglia*

| Comparison (Treatment) | ANOVA p-value |
|---|---|
| MTZ vs DMSO | 8.9e-06 |

| Comparison (Genotype) | ANOVA p-value |
|---|---|
| *w211+/-* vs *w211-/-* | 0.3640735 |

| Comparison | post-hoc Tukey's HSD |
|---|---|
| DMSO *w211 +/-* vs MTZ *w211 +/-* | 0.0025799 |
| DMSO *w211 +/-* vs DMSO *w211 -/-* | 0.7956819 |
| DMSO *w211 +/-* vs MTZ *w211 -/-* | 0.0060499 |
| DMSO *w211 -/-* vs MTZ *w211 +/-* | 0.0011806 |
| MTZ *w211 -/-* vs MTZ *w211 +/-* | 0.9638798 |
| DMSO *w211 -/-* vs MTZ *w211 -/-* | 0.0024917 |

### Figure 6:

*Microglia*

| Comparison (Treatment) | ANOVA p-value |
|---|---|
| MTZ vs DMSO | 0.0005235 |

| Comparison (Genotype) | ANOVA p-value |
|---|---|
| *w211+/-* vs *w211-/-* | 0.8666362 |

| Comparison | post-hoc Tukey's HSD |
|---|---|
| DMSO *w211 +/-* vs MTZ *w211 +/-* | 0.0066590 |
| DMSO *w211 +/-* vs DMSO *w211 -/-* | 0.9087570 |
| DMSO *w211 +/-* vs MTZ *w211 -/-* | 0.0527973 |
| DMSO *w211 -/-* vs MTZ *w211 +/-* | 0.0431927 |
| MTZ *w211 -/-* vs MTZ *w211 +/-* | 0.8152699 |
| DMSO *w211 -/-* vs MTZ *w211 -/-* | 0.2305585 |

*Activated morphology*

| Comparison (Treatment) | ANOVA p-value |
|---|---|
| MTZ vs DMSO | 0.4962307 |

| Comparison (Genotype) | ANOVA p-value |
|---|---|
| *w211+/-* vs *w211-/-* | 0.1635403 |

| Comparison | post-hoc Tukey's HSD |
|---|---|
| DMSO *w211 +/-* vs MTZ *w211 +/-* | 0.9985847 |
| DMSO *w211 +/-* vs DMSO *w211 -/-* | 0.9842636 |
| DMSO *w211 +/-* vs MTZ *w211 -/-* | 0.4559497 |
| DMSO *w211 -/-* vs MTZ *w211 +/-* | 0.9566604 |
| MTZ *w211 -/-* vs MTZ *w211 +/-* | 0.3693206 |
| DMSO *w211 -/-* vs MTZ *w211 -/-* | 0.7008686 |

*ONL/OPL-localized microglia*

| Comparison (Treatment) | ANOVA p-value |
|---|---|
| MTZ vs DMSO | 0.0163176 |

| Comparison (Genotype) | ANOVA p-value |
|---|---|
| *w211+/-* vs *w211-/-* | 0.6369304 |

| Comparison | post-hoc Tukey's HSD |
|---|---|
| DMSO *w211 +/-* vs MTZ *w211 +/-* | 0.1391822 |
| DMSO *w211 +/-* vs DMSO *w211 -/-* | 0.8749932 |
| DMSO *w211 +/-* vs MTZ *w211 -/-* | 0.1563744 |
| DMSO *w211 -/-* vs MTZ *w211 +/-* | 0.5054537 |

| | |
|---|---|
| MTZ *w211 -/-* vs MTZ *w211 +/-* | 0.9999105 |
| DMSO *w211 -/-* vs MTZ *w211 -/-* | 0.5417926 |

*Total TUNEL+ cells*

| Comparison (Treatment) | ANOVA p-value |
|---|---|
| MTZ vs DMSO | 0.4771231 |

| Comparison (Genotype) | ANOVA p-value |
|---|---|
| *w211+/-* vs *w211-/-* | 0.0051213 |

| Comparison | post-hoc Tukey's HSD |
|---|---|
| DMSO *w211 +/-* vs MTZ *w211 +/-* | 0.8596009 |
| DMSO *w211 +/-* vs DMSO *w211 -/-* | 0.0874449 |
| DMSO *w211 +/-* vs MTZ *w211 -/-* | 0.0619094 |
| DMSO *w211 -/-* vs MTZ *w211 +/-* | 0.3374423 |
| MTZ *w211 -/-* vs MTZ *w211 +/-* | 0.2727643 |
| DMSO *w211 -/-* vs MTZ *w211 -/-* | 0.9998384 |

*ONL TUNEL+ cells*

Conover test with Bonferroni correction

| Comparison | p-value |
|---|---|
| DMSO *w211 +/-* vs MTZ *w211 +/-* | 0.0035 |
| DMSO *w211 +/-* vs DMSO *w211 -/-* | 1.0000 |
| DMSO *w211 +/-* vs MTZ *w211 -/-* | 0.0207 |
| DMSO *w211 -/-* vs MTZ *w211 +/-* | 0.0125 |
| MTZ *w211 -/-* vs MTZ *w211 +/-* | 1.0000 |
| DMSO *w211 -/-* vs MTZ *w211 -/-* | 0.0632 |

*Apoptotic Microglia*

| Comparison | post-hoc Tukey's HSD |
|---|---|
| DMSO *w211 +/-* vs MTZ *w211 +/-* | 1.0000 |
| DMSO *w211 +/-* vs DMSO *w211 -/-* | 0.0086 |
| DMSO *w211 +/-* vs MTZ *w211 -/-* | 1.0000 |
| DMSO *w211 -/-* vs MTZ *w211 +/-* | 0.0086 |
| MTZ *w211 -/-* vs MTZ *w211 +/-* | 1.0000 |
| DMSO *w211 -/-* vs MTZ *w211 -/-* | 0.0086 |

### Figure 7:

*48 hpt*

Total PCNA

| Comparison (Treatment) | ANOVA p-value |
|---|---|
| MTZ vs DMSO | 0.000813 |

| Comparison (Genotype) | ANOVA p-value |
|---|---|
| *w211+/-* vs *w211-/-* | 0.5889081 |

| Comparison | post-hoc Tukey's HSD |
|---|---|
| DMSO *w211 +/-* vs MTZ *w211 +/-* | 0.0988665 |
| DMSO *w211 +/-* vs DMSO *w211 -/-* | 0.9877349 |
| DMSO *w211 +/-* vs MTZ *w211 -/-* | 0.0880629 |
| DMSO *w211 -/-* vs MTZ *w211 +/-* | 0.0396134 |
| MTZ *w211 -/-* vs MTZ *w211 +/-* | 0.9673937 |
| DMSO *w211 -/-* vs MTZ *w211 -/-* | 0.0251719 |

ONL PCNA

| Comparison (Treatment) | ANOVA p-value |
|---|---|
| MTZ vs DMSO | 0.0017031 |

| Comparison (Genotype) | ANOVA p-value |
|---|---|
| *w211+/-* vs *w211-/-* | 0.9796476 |

| Comparison | post-hoc Tukey's HSD |
|---|---|
| DMSO *w211 +/-* vs MTZ *w211 +/-* | 0.9740744 |
| DMSO *w211 +/-* vs DMSO *w211 -/-* | 0.3656315 |
| DMSO *w211 +/-* vs MTZ *w211 -/-* | 0.2483564 |
| DMSO *w211 -/-* vs MTZ *w211 +/-* | 0.1831137 |
| MTZ *w211 -/-* vs MTZ *w211 +/-* | 0.5037547 |
| DMSO *w211 -/-* vs MTZ *w211 -/-* | 0.0025015 |

INL PCNA

| Comparison (Treatment) | ANOVA p-value |
|---|---|
| MTZ vs DMSO | 0.004012 |

| Comparison (Genotype) | ANOVA p-value |
|---|---|
| *w211+/-* vs *w211-/-* | 0.343975 |

| Comparison | post-hoc Tukey's HSD |
|---|---|
| DMSO *w211 +/-* vs MTZ *w211 +/-* | 0.0424281 |
| DMSO *w211 +/-* vs DMSO *w211 -/-* | 0.9989466 |
| DMSO *w211 +/-* vs MTZ *w211 -/-* | 0.2016223 |
| DMSO *w211 -/-* vs MTZ *w211 +/-* | 0.0412237 |
| MTZ *w211 -/-* vs MTZ *w211 +/-* | 0.4959494 |
| DMSO *w211 -/-* vs MTZ *w211 -/-* | 0.2038981 |

PCNA+ Müller glia cells

| Comparison (Treatment) | ANOVA p-value |
|---|---|
| MTZ vs DMSO | 0.0005101 |

| Comparison (Genotype) | ANOVA p-value |
|---|---|
| *w211*+/- vs *w211*-/- | 0.5005205 |

| Comparison | post-hoc Tukey's HSD |
|---|---|
| DMSO *w211* +/- vs MTZ *w211* +/- | 0.0133734 |
| DMSO *w211* +/- vs DMSO *w211* -/- | 0.9822362 |
| DMSO *w211* +/- vs MTZ *w211* -/- | 0.0413399 |
| DMSO *w211* -/- vs MTZ *w211* +/- | 0.0200409 |
| MTZ *w211* -/- vs MTZ *w211* +/- | 0.6055039 |
| DMSO *w211* -/- vs MTZ *w211* -/- | 0.0648688 |

Rod cells

| Comparison (Treatment) | ANOVA p-value |
|---|---|
| MTZ vs DMSO | 8.9e-06 |

| Comparison (Genotype) | ANOVA p-value |
|---|---|
| *w211*+/- vs *w211*-/- | 0.5909452 |

| Comparison | post-hoc Tukey's HSD |
|---|---|
| DMSO *w211* +/- vs MTZ *w211* +/- | 0.0091936 |
| DMSO *w211* +/- vs DMSO *w211* -/- | 0.9615604 |
| DMSO *w211* +/- vs MTZ *w211* -/- | 0.0008076 |
| DMSO *w211* -/- vs MTZ *w211* +/- | 0.0174790 |
| MTZ *w211* -/- vs MTZ *w211* +/- | 0.9888983 |

| | |
|---|---|
| DMSO *w211 -/-* vs MTZ *w211 -/-* | 0.0013304 |

*72 hpt*

Total PCNA

| Comparison (Treatment) | ANOVA p-value |
|---|---|
| MTZ vs DMSO | 7e-07 |

| Comparison (Genotype) | ANOVA p-value |
|---|---|
| *w211+/-* vs *w211-/-* | 0.6504237 |

| Comparison | post-hoc Tukey's HSD |
|---|---|
| DMSO *w211 +/-* vs MTZ *w211 +/-* | 0.0000821 |
| DMSO *w211 +/-* vs DMSO *w211 -/-* | 0.9969671 |
| DMSO *w211 +/-* vs MTZ *w211 -/-* | 0.0014340 |
| DMSO *w211 -/-* vs MTZ *w211 +/-* | 0.0003291 |
| MTZ *w211 -/-* vs MTZ *w211 +/-* | 0.8757365 |
| DMSO *w211 -/-* vs MTZ *w211 -/-* | 0.0041165 |

ONL PCNA

| Comparison (Treatment) | ANOVA p-value |
|---|---|
| MTZ vs DMSO | 0.0155224 |

| Comparison (Genotype) | ANOVA p-value |
|---|---|
| *w211+/-* vs *w211-/-* | 0.0455237 |

| Comparison | post-hoc Tukey's HSD |
|---|---|

| | |
|---|---|
| DMSO *w211 +/-* vs MTZ *w211 +/-* | 0.0107579 |
| DMSO *w211 +/-* vs DMSO *w211 -/-* | 0.9442936 |
| DMSO *w211 +/-* vs MTZ *w211 -/-* | 0.9790447 |
| DMSO *w211 -/-* vs MTZ *w211 +/-* | 0.0689834 |
| MTZ *w211 -/-* vs MTZ *w211 +/-* | 0.0208466 |
| DMSO *w211 -/-* vs MTZ *w211 -/-* | 0.9970386 |

INL PCNA

| Comparison (Treatment) | ANOVA p-value |
|---|---|
| MTZ vs DMSO | 6e-07 |

| Comparison (Genotype) | ANOVA p-value |
|---|---|
| *w211+/-* vs *w211-/-* | 0.4259122 |

| Comparison | post-hoc Tukey's HSD |
|---|---|
| DMSO *w211 +/-* vs MTZ *w211 +/-* | 0.0001927 |
| DMSO *w211 +/-* vs DMSO *w211 -/-* | 0.9549493 |
| DMSO *w211 +/-* vs MTZ *w211 -/-* | 0.0001477 |
| DMSO *w211 -/-* vs MTZ *w211 +/-* | 0.0017826 |
| MTZ *w211 -/-* vs MTZ *w211 +/-* | 0.9222452 |
| DMSO *w211 -/-* vs MTZ *w211 -/-* | 0.0011009 |

PCNA+ Müller glia cells

| Comparison (Treatment) | ANOVA p-value |
|---|---|
| MTZ vs DMSO | 0.0698587 |

| Comparison (Genotype) | ANOVA p-value |
|---|---|
| *w211+/-* vs *w211-/-* | 0.0700307 |

| Comparison | post-hoc Tukey's HSD |
|---|---|
| DMSO *w211 +/-* vs MTZ *w211 +/-* | 0.0466808 |
| DMSO *w211 +/-* vs DMSO *w211 -/-* | 0.9475726 |
| DMSO *w211 +/-* vs MTZ *w211 -/-* | 0.9999475 |
| DMSO *w211 -/-* vs MTZ *w211 +/-* | 0.2117242 |
| MTZ *w211 -/-* vs MTZ *w211 +/-* | 0.0393548 |
| DMSO *w211 -/-* vs MTZ *w211 -/-* | 0.9554775 |

Rod cells

| Comparison (Treatment) | ANOVA p-value |
|---|---|
| MTZ vs DMSO | 1e-07 |

| Comparison (Genotype) | ANOVA p-value |
|---|---|
| *w211+/-* vs *w211-/-* | 0.1884319 |

| Comparison | post-hoc Tukey's HSD |
|---|---|
| DMSO *w211 +/-* vs MTZ *w211 +/-* | 0.0000005 |
| DMSO *w211 +/-* vs DMSO *w211 -/-* | 0.5798035 |
| DMSO *w211 +/-* vs MTZ *w211 -/-* | 0.0009044 |
| DMSO *w211 -/-* vs MTZ *w211 +/-* | 0.0000458 |
| MTZ *w211 -/-* vs MTZ *w211 +/-* | 0.0488950 |
| DMSO *w211 -/-* vs MTZ *w211 -/-* | 0.0385270 |

*96 hpt*

Total PCNA

| Comparison (Treatment) | ANOVA p-value |
|---|---|
| MTZ vs DMSO | 3e-07 |

| Comparison (Genotype) | ANOVA p-value |
|---|---|
| *w211*+/- vs *w211*-/- | 0.8562333 |

| Comparison | post-hoc Tukey's HSD |
|---|---|
| DMSO *w211* +/- vs MTZ *w211* +/- | 0.0002655 |
| DMSO *w211* +/- vs DMSO *w211* -/- | 0.9988123 |
| DMSO *w211* +/- vs MTZ *w211* -/- | 0.0008579 |
| DMSO *w211* -/- vs MTZ *w211* +/- | 0.0000936 |
| MTZ *w211* -/- vs MTZ *w211* +/- | 0.9994374 |
| DMSO *w211* -/- vs MTZ *w211* -/- | 0.0003602 |

ONL PCNA

| Comparison (Treatment) | ANOVA p-value |
|---|---|
| MTZ vs DMSO | 0.2818261 |

| Comparison (Genotype) | ANOVA p-value |
|---|---|
| *w211*+/- vs *w211*-/- | 0.549114 |

| Comparison | post-hoc Tukey's HSD |
|---|---|
| DMSO *w211* +/- vs MTZ *w211* +/- | 0.9568367 |
| DMSO *w211* +/- vs DMSO *w211* -/- | 0.3260329 |
| DMSO *w211* +/- vs MTZ *w211* -/- | 0.9982579 |
| DMSO *w211* -/- vs MTZ *w211* +/- | 0.4929352 |
| MTZ *w211* -/- vs MTZ *w211* +/- | 0.8879518 |
| DMSO *w211* -/- vs MTZ *w211* -/- | 0.2203008 |

INL PCNA

| Comparison (Treatment) | ANOVA p-value |
|---|---|
| MTZ vs DMSO | 2e-07 |

| Comparison (Genotype) | ANOVA p-value |
|---|---|
| *w211+/-* vs *w211-/-* | 0.9292638 |

| Comparison | post-hoc Tukey's HSD |
|---|---|
| DMSO *w211 +/-* vs MTZ *w211 +/-* | 0.0000553 |
| DMSO *w211 +/-* vs DMSO *w211 -/-* | 0.9583070 |
| DMSO *w211 +/-* vs MTZ *w211 -/-* | 0.0005639 |
| DMSO *w211 -/-* vs MTZ *w211 +/-* | 0.0001217 |
| MTZ *w211 -/-* vs MTZ *w211 +/-* | 0.9419078 |
| DMSO *w211 -/-* vs MTZ *w211 -/-* | 0.0013362 |

PCNA+ Müller glia cells

| Comparison (Treatment) | ANOVA p-value |
|---|---|
| MTZ vs DMSO | 0.2188456 |

| Comparison (Genotype) | ANOVA p-value |
|---|---|
| *w211+/-* vs *w211-/-* | 0.3782837 |

| Comparison | post-hoc Tukey's HSD |
|---|---|
| DMSO *w211 +/-* vs MTZ *w211 +/-* | 0.7934404 |
| DMSO *w211 +/-* vs DMSO *w211 -/-* | 0.4804981 |
| DMSO *w211 +/-* vs MTZ *w211 -/-* | 0.5396239 |
| DMSO *w211 -/-* vs MTZ *w211 +/-* | 0.9067730 |
| MTZ *w211 -/-* vs MTZ *w211 +/-* | 0.0848247 |

| | |
|---|---|
| DMSO *w211 -/-* vs MTZ *w211 -/-* | 0.0340715 |

*Rod cells*

| Comparison (Treatment) | ANOVA p-value |
|---|---|
| MTZ vs DMSO | 2e-07 |

| Comparison (Genotype) | ANOVA p-value |
|---|---|
| *w211+/-* vs *w211-/-* | 0.4529395 |

| Comparison | post-hoc Tukey's HSD |
|---|---|
| DMSO *w211 +/-* vs MTZ *w211 +/-* | 0.0004111 |
| DMSO *w211 +/-* vs DMSO *w211 -/-* | 0.8865608 |
| DMSO *w211 +/-* vs MTZ *w211 -/-* | 0.0023621 |
| DMSO *w211 -/-* vs MTZ *w211 +/-* | 0.0000274 |
| MTZ *w211 -/-* vs MTZ *w211 +/-* | 0.9802286 |
| DMSO *w211 -/-* vs MTZ *w211 -/-* | 0.0002202 |

*120 hpt*

*Total PCNA*

| Comparison (Treatment) | ANOVA p-value |
|---|---|
| MTZ vs DMSO | 1.4e-06 |

| Comparison (Genotype) | ANOVA p-value |
|---|---|
| *w211+/-* vs *w211-/-* | 0.7549218 |

| Comparison | post-hoc Tukey's HSD |
|---|---|
| DMSO *w211 +/-* vs MTZ *w211 +/-* | 0.0001536 |

| | |
|---|---|
| DMSO *w211 +/-* vs DMSO *w211 -/-* | 0.9987829 |
| DMSO *w211 +/-* vs MTZ *w211 -/-* | 0.0015040 |
| DMSO *w211 -/-* vs MTZ *w211 +/-* | 0.0008546 |
| MTZ *w211 -/-* vs MTZ *w211 +/-* | 0.9488673 |
| DMSO *w211 -/-* vs MTZ *w211 -/-* | 0.0054610 |

*ONL PCNA*

| Comparison (Treatment) | ANOVA p-value |
|---|---|
| MTZ vs DMSO | 0.6183428 |

| Comparison (Genotype) | ANOVA p-value |
|---|---|
| *w211+/-* vs *w211-/-* | 0.1321385 |

| Comparison | post-hoc Tukey's HSD |
|---|---|
| DMSO *w211 +/-* vs MTZ *w211 +/-* | 0.9997994 |
| DMSO *w211 +/-* vs DMSO *w211 -/-* | 0.9709647 |
| DMSO *w211 +/-* vs MTZ *w211 -/-* | 0.4760440 |
| DMSO *w211 -/-* vs MTZ *w211 +/-* | 0.9474022 |
| MTZ *w211 -/-* vs MTZ *w211 +/-* | 0.3628292 |
| DMSO *w211 -/-* vs MTZ *w211 -/-* | 0.8088739 |

*INL PCNA*

| Comparison (Treatment) | ANOVA p-value |
|---|---|
| MTZ vs DMSO | 1.3e-06 |

| Comparison (Genotype) | ANOVA p-value |
|---|---|
| *w211+/-* vs *w211-/-* | 0.8376779 |

| Comparison | post-hoc Tukey’s HSD |
|---|---|
| DMSO *w211* +/- vs MTZ *w211* +/- | 0.0000436 |
| DMSO *w211* +/- vs DMSO *w211* -/- | 0.7169200 |
| DMSO *w211* +/- vs MTZ *w211* -/- | 0.0005727 |
| DMSO *w211* -/- vs MTZ *w211* +/- | 0.0032287 |
| MTZ *w211* -/- vs MTZ *w211* +/- | 0.9244720 |
| DMSO *w211* -/- vs MTZ *w211* -/- | 0.0209319 |

*PCNA+ Müller glia cells*

| Comparison (Treatment) | ANOVA p-value |
|---|---|
| MTZ vs DMSO | 0.6729541 |

| Comparison (Genotype) | ANOVA p-value |
|---|---|
| *w211+/-* vs *w211-/-* | 0.3117225 |

| Comparison | post-hoc Tukey’s HSD |
|---|---|
| DMSO *w211* +/- vs MTZ *w211* +/- | 0.9293720 |
| DMSO *w211* +/- vs DMSO *w211* -/- | 0.9874013 |
| DMSO *w211* +/- vs MTZ *w211* -/- | 0.9726581 |
| DMSO *w211* -/- vs MTZ *w211* +/- | 0.8066538 |
| MTZ *w211* -/- vs MTZ *w211* +/- | 0.7090040 |
| DMSO *w211* -/- vs MTZ *w211* -/- | 0.9999032 |

*Rod cells*

| Comparison (Treatment) | ANOVA p-value |
|---|---|
| MTZ vs DMSO | 7e-07 |

| Comparison (Genotype) | ANOVA p-value |
|---|---|
| *w211+/-* vs *w211-/-* | 0.8087558 |

| Comparison | post-hoc Tukey's HSD |
|---|---|
| DMSO *w211 +/-* vs MTZ *w211 +/-* | 0.0000270 |
| DMSO *w211 +/-* vs DMSO *w211 -/-* | 0.7132361 |
| DMSO *w211 +/-* vs MTZ *w211 -/-* | 0.0003306 |
| DMSO *w211 -/-* vs MTZ *w211 +/-* | 0.0021432 |
| MTZ *w211 -/-* vs MTZ *w211 +/-* | 0.9380778 |
| DMSO *w211 -/-* vs MTZ *w211 -/-* | 0.0134222 |

### Supplemental Figure 3:

*p62*

| Comparison (Treatment) | ANOVA p-value |
|---|---|
| MTZ vs DMSO | 0.0003572 |

| Comparison (Genotype) | ANOVA p-value |
|---|---|
| *w211+/-* vs *w211-/-* | 0.4491311 |

| Comparison | post-hoc Tukey's HSD |
|---|---|
| DMSO *w211 +/-* vs MTZ *w211 +/-* | 0.0052249 |
| DMSO *w211 +/-* vs DMSO *w211 -/-* | 0.9952397 |
| DMSO *w211 +/-* vs MTZ *w211 -/-* | 0.0492740 |
| DMSO *w211 -/-* vs MTZ *w211 +/-* | 0.0111252 |

| | |
|---|---|
| MTZ *w211 -/-* vs MTZ *w211 +/-* | 0.5189272 |
| DMSO *w211 -/-* vs MTZ *w211 -/-* | 0.0990189 |

*gpx1a*

| Comparison (Treatment) | ANOVA p-value |
|---|---|
| MTZ vs DMSO | 0.9108778 |

| Comparison (Genotype) | ANOVA p-value |
|---|---|
| *w211+/-* vs *w211-/-* | 0.3016868 |

| Comparison | post-hoc Tukey's HSD |
|---|---|
| DMSO *w211 +/-* vs MTZ *w211 +/-* | 0.9983820 |
| DMSO *w211 +/-* vs DMSO *w211 -/-* | 0.8513942 |
| DMSO *w211 +/-* vs MTZ *w211 -/-* | 0.9249873 |
| DMSO *w211 -/-* vs MTZ *w211 +/-* | 0.8133448 |
| MTZ *w211 -/-* vs MTZ *w211 +/-* | 0.8893597 |
| DMSO *w211 -/-* vs MTZ *w211 -/-* | 0.9963259 |

*nfe2l2a*

| Comparison (Treatment) | ANOVA p-value |
|---|---|
| MTZ vs DMSO | 0.1192503 |

| Comparison (Genotype) | ANOVA p-value |
|---|---|
| *w211+/-* vs *w211-/-* | 0.9152659 |

| Comparison | post-hoc Tukey's HSD |
|---|---|
| DMSO *w211 +/-* vs MTZ *w211 +/-* | 0.9534378 |
| DMSO *w211 +/-* vs DMSO *w211 -/-* | 0.9158634 |

| | |
|---|---|
| DMSO *w211 +/-* vs MTZ *w211 -/-* | 0.6528431 |
| DMSO *w211 -/-* vs MTZ *w211 +/-* | 0.7051852 |
| MTZ *w211 -/-* vs MTZ *w211 +/-* | 0.9511098 |
| DMSO *w211 -/-* vs MTZ *w211 -/-* | 0.3250577 |

*nfe2l2b*

| Comparison (Treatment) | ANOVA p-value |
|---|---|
| MTZ vs DMSO | 0.0658942 |

| Comparison (Genotype) | ANOVA p-value |
|---|---|
| *w211+/-* vs *w211-/-* | 0.699699 |

| Comparison | post-hoc Tukey's HSD |
|---|---|
| DMSO *w211 +/-* vs MTZ *w211 +/-* | 0.6256223 |
| DMSO *w211 +/-* vs DMSO *w211 -/-* | 0.9961647 |
| DMSO *w211 +/-* vs MTZ *w211 -/-* | 0.3315585 |
| DMSO *w211 -/-* vs MTZ *w211 +/-* | 0.7665372 |
| MTZ *w211 -/-* vs MTZ *w211 +/-* | 0.9852050 |
| DMSO *w211 -/-* vs MTZ *w211 -/-* | 0.4891223 |

*il1b*

| Comparison (Treatment) | ANOVA p-value |
|---|---|
| MTZ vs DMSO | 0.4280449 |

| Comparison (Genotype) | ANOVA p-value |
|---|---|
| *w211+/-* vs *w211-/-* | 0.3996548 |

| Comparison | post-hoc Tukey's HSD |
|---|---|

| | |
|---|---|
| DMSO *w211 +/-* vs MTZ *w211 +/-* | 0.6576556 |
| DMSO *w211 +/-* vs DMSO *w211 -/-* | 0.5794538 |
| DMSO *w211 +/-* vs MTZ *w211 -/-* | 0.6438940 |
| DMSO *w211 -/-* vs MTZ *w211 +/-* | 0.9999563 |
| MTZ *w211 -/-* vs MTZ *w211 +/-* | 0.9996039 |
| DMSO *w211 -/-* vs MTZ *w211 -/-* | 0.9983990 |

*bip*

| Comparison (Treatment) | ANOVA p-value |
|---|---|
| MTZ vs DMSO | 0.6070691 |

| Comparison (Genotype) | ANOVA p-value |
|---|---|
| *w211+/-* vs *w211-/-* | 0.4954893 |

| Comparison | post-hoc Tukey's HSD |
|---|---|
| DMSO *w211 +/-* vs MTZ *w211 +/-* | 0.9968102 |
| DMSO *w211 +/-* vs DMSO *w211 -/-* | 0.6915930 |
| DMSO *w211 +/-* vs MTZ *w211 -/-* | 0.9999034 |
| DMSO *w211 -/-* vs MTZ *w211 +/-* | 0.8537129 |
| MTZ *w211 -/-* vs MTZ *w211 +/-* | 0.9987851 |
| DMSO *w211 -/-* vs MTZ *w211 -/-* | 0.7266996 |

*gfap*

| Comparison (Treatment) | ANOVA p-value |
|---|---|
| MTZ vs DMSO | 0.8610488 |

| Comparison (Genotype) | ANOVA p-value |
|---|---|
| *w211+/-* vs *w211-/-* | 0.8643464 |

| Comparison | post-hoc Tukey's HSD |
|---|---|
| DMSO *w211 +/-* vs MTZ *w211 +/-* | 0.9937994 |
| DMSO *w211 +/-* vs DMSO *w211 -/-* | 0.9926402 |
| DMSO *w211 +/-* vs MTZ *w211 -/-* | 0.9943641 |
| DMSO *w211 -/-* vs MTZ *w211 +/-* | 1.0000000 |
| MTZ *w211 -/-* vs MTZ *w211 +/-* | 0.9999846 |
| DMSO *w211 -/-* vs MTZ *w211 -/-* | 0.9999836 |

*vim*

| Comparison (Treatment) | ANOVA p-value |
|---|---|
| MTZ vs DMSO | 0.1745582 |

| Comparison (Genotype) | ANOVA p-value |
|---|---|
| *w211+/-* vs *w211-/-* | 0.1647218 |

| Comparison | post-hoc Tukey's HSD |
|---|---|
| DMSO *w211 +/-* vs MTZ *w211 +/-* | 0.9912232 |
| DMSO *w211 +/-* vs DMSO *w211 -/-* | 0.1193918 |
| DMSO *w211 +/-* vs MTZ *w211 -/-* | 0.9994615 |
| DMSO *w211 -/-* vs MTZ *w211 +/-* | 0.2776078 |
| MTZ *w211 -/-* vs MTZ *w211 +/-* | 0.9790807 |
| DMSO *w211 -/-* vs MTZ *w211 -/-* | 0.0982257 |
